# Detection of bacteria through taste receptors primes the cellular immune response

**DOI:** 10.1101/2024.09.26.615243

**Authors:** Alix Najera Mazariegos, Gérard Manière, Léo Sillon, Romane Milleville, Martine Berthelot-Grosjean, Enisa Aruçi, Darius Camp, George Alves, Rhea Kaul, Carla Jane Duval, Isabelle Chauvel, Julien Royet, Yaël Grosjean, Pierre-Yves Musso, Guy Tanentzapf

## Abstract

Animals use their sensory system to detect cues in their external environment, then communicate, process, and integrate these cues through the nervous system in order to elicit a specific response. Taste is an important cue used by animals to explore their external environment and can modulate various aspects of animal behavior and physiology. A major ongoing challenge for animals is to detect and respond to the presence of a variety of microbes in their environment. However, to date, the links between the sensory system and the response to pathogenic threats remain poorly understood. Here we show that *Drosophila melanogaster* larvae use their taste system to detect bacterial peptidoglycans in their environment and respond by modulating the activity of their cellular immune system. We show that specific PeptidoGlycan Receptor Proteins (PGRPs) act in aversive taste neurons, via a non-canonical Immune Deficiency (Imd) pathway. These PGRPs mediate signaling in taste neurons and control immune cells production in the larval hematopoietic organ, the lymph gland. Taste-mediated sensing of bacteria in larvae primes the immune system, and improves survival after infection in adult flies. These results demonstrate that sensory inputs such as taste play an important role in protecting animals from bacterial infection by providing a powerful adaptive response to potential pathogens. Overall, our findings add to the growing list of examples of crosstalk between the nervous and immune systems and provide novel and important mechanisms for linking them.

**One Sentence Summary:** Najera Mazariegos et al. demonstrate that organisms can use taste to monitor their environment for potential immune challenges and activate their immune system if they detect bacteria.

## INTRODUCTION

Animal homeostasis requires constant monitoring of the environment so that animals can adapt and respond to external cues. Key to animal survival is their ability to detect the presence of both the nutrients they need to survive and possible pathogens and toxins in the environment. The detection and analysis of chemical cues through the senses of smell and taste is an important method used by animals to monitor their environment. Smell and taste are used by animals to communicate, detect food, analyze food quality, and drive ingestion. In humans, for example, the detection of food decomposition through taste and smell drives avoidance or rejection behavior (Wisman and Shrira, 2015), whereas food-associated odor and taste drive attraction towards the food source (Boesveldt and Graaf, 2017). Although such behaviors vary widely among animals and are context specific, the general regulatory principles are conserved. This is exemplified by taste modalities that are well conserved in animals ranging from humans to *Drosophila melanogaster* (Wisman and Shrira, 2015).

After ingestion, food is digested and evaluated through post-ingestive processes (Musso et al., 2017; Sclafani, 2001). High energy content will assign a positive value to the food, which is regulated by neurohormonal pathways such as insulin signaling (Ratnaparkhi and Sudhakaran, 2022); whereas a post-ingestive illness will assign a negative value to the food, and lead to a biological response aimed at reestablishing homeostasis (Lin et al., 2018; Wright et al., 2010; Yao and Scott, 2022). In particular, toxic food-borne compounds can induce a stereotyped immune response to restore homeostasis. Pathogen recognition can trigger a variety of immune-mediated responses to either remove or destroy the threat (Yu et al., 2022). Although immune responses are generally the result of an infection, *i.e.* the entry of pathogens into the organism, pathogen-associated olfactory cues can act at a distance to control infection-avoidance behavior, without direct contact between the animal and the pathogen (Shim et al., 2013). However, little is known about whether or how the detection of sensory cues can indicate a pathogenic threat, prime the immune response, or prepare the organism for the possibility of an infectious challenge.

Drosophila provides a powerful genetically tractable model system for studying immunity. Similar to mammals, flies possess both humoral and cellular immune responses (Yu et al., 2022). Humoral immunity involves the regulated expression of genes encoding antimicrobial peptides (AMPs) that disrupt membranes and internal functions of pathogens in order to kill them. On the other hand, in invertebrates, cellular immunity involves recruitment of specialized immune blood cells, called hemocytes. Hemocytes are recruited to the site of infection and kill pathogens directly by phagocytosis. These cellular immune responses are activated immediately upon pathogen detection and substantially limit microbial proliferation (Yu et al., 2022).

In Drosophila larvae, immune cells are produced in a specialized organ called the **<u>L</u>**ymph **<u>G</u>**land (LG), located anterior to the heart tube. Several factors make the *Drosophila* larvae a useful model for studying hematopoiesis, including its mechanistic conservation with vertebrate blood development, the wide availability of genetic tools and reagents, and its accessibility for imaging (Banerjee et al., 2019). During fly hematopoiesis, a distinct population of blood progenitors gives rise to blood cells that serve as the fly’s cellular immune system. Drosophila blood progenitors, known as prohemocytes, exhibit many stem-cell-like properties and their behavior is controlled by a small population of specialized cells that act as a hematopoietic niche, the Posterior Signaling Centre (PSC) (Cho et al., 2020; Krzemień et al., 2007; Mandal et al., 2007; Tokusumi et al., 2010). Prohemocytes can give rise to plasmatocytes, lamellocytes, and crystal cells, three highly differentiated blood cell types that have unique functions and roles in the cellular immune response in flies (Jung et al., 2005). Specifically, plasmatocytes, that represent more than 90% of hemocytes, are the functional equivalent of human macrophages and engulf and consume microbes and dead host cells; crystal cells drive melanization for wound repair; while lamellocytes are expressed only in response to parasitic wasp infestation. The *Drosophila* immune system is activated by direct contact of immune cells with bacterial pathogens. This response is driven by toll-like and peptidoglycan receptors that recognize a variety of bacteria by direct binding but can also be modulated by environmental cues (Bland, 2023; Lu et al., 2020; Shim et al., 2013; Yu et al., 2022). For example, anosmic flies, which are unable to smell, exhibit increased blood cell differentiation, suggesting that chemically based sensory inputs may regulate the cellular immune response (Shim et al., 2013).

There is considerable conservation in how animals detect and respond to taste. Sensory cells in both flies and mammals express gustatory receptors that are specialized in the detection of specific tastants, the activation of which drives specific behaviors, exemplified by the appetitive response to sweet taste and the aversive response to bitter taste (Yarmolinsky et al., 2009). Taste modalities are generally divided into 5 families: Sweet, Bitter, Umami, Sour, and Salt. However, the ongoing discovery of new taste sensory pathways such as bacteria sensing or fatty acid sensing suggest an even greater complexity to the sense of taste. There are multiple examples of taste controlling behavior in flies. For example, taste-mediated detection of the chemical signature of Gram-negative bacteria in adult flies, and subsequent activation of the immune-specific Imd pathway in the taste sensory organs, induces a refusal to lay eggs in contaminated areas and a rejection of contaminated-food (Masuzzo et al., 2022; Montanari et al., 2024). Another example of the complex behavior controlled by taste modalities in flies is the observation that sweet taste acts as a predictor of metabolic value and consequently regulates feeding (Dus et al., 2013, 2011; Musso et al., 2017, 2015; Yao and Scott, 2022). The question arises as to whether the pathways that act downstream of taste go beyond modifying behavior and alter other aspects of animal homeostasis in response to the detection of the chemical signature of potentially pathogenic bacteria.

While sensory detection of pathogens is known to drive behavioural avoidance, it remains unclear whether sensory cues can also actively shape internal immune pathways before infection occurs. Recent work has shown that olfactory cues linked to parasite exposure can promote lamellocyte differentiation in anticipation of wasp infection (Madhwal et al., 2020; Shim et al., 2013), raising the possibility that sensory systems may not only detect threats but also prepare immune defenses. Because taste neurons of adult Drosophila can detect bacteria-derived peptidoglycans (PGNs) via the PGRP-LC receptor (Masuzzo et al., 2022; Soldano et al., 2016) and send protection to the brain, they are well positioned to serve as neural immune sensors. However, whether taste circuits use bacterial cues to modulate hematopoiesis, rather than just behavioural avoidance, has not been explored.

Here we show that, in *Drosophila*, the detection of bacteria-derived pathogen-associated molecular patterns (PAMPs) via taste regulates homeostasis in an unexpected way, by modulating hematopoiesis. Specifically, we show that the genetic silencing of specific aversive bitter taste-sensing peripheral neurons increases plasmatocyte production in the LG. RNAi-mediated gene knockdown shows that the well-established bitter taste receptor, Gr66a, does not alter immune homeostasis. However, we find that aversive taste neurons detect Gram-negative bacteria via the peptidoglycan receptor PGRP-LC, which then activates a non-canonical Imd pathway to suppress neural activity that promotes plasmatocyte differentiation in the LG. In addition, suppressing detection of Gram-negative bacteria reduces plasmatocyte differentiation, while overactivation of the Imd pathway within aversive taste neurons strongly promotes plasmatocyte differentiation and improves survival to infection in adult flies. This demonstrates that the sense of taste plays a role in the homeostatic regulation of immunity and its adaptation to pathogens throughout life.

## RESULTS

### Drosophila larvae detect Gram-negative bacteria-derived peptidoglycans through Gr66a taste neurons

To date the question of whether fly larva can detect bacteria via taste has not been addressed. It is known however that in the adult fly, a subpopulation of taste-sensing neurons, characterized by the expression of the gustatory receptor Gr66a, is capable of sensing bacteria-derived peptidoglycans (PGN) (Masuzzo et al., 2022). Specifically, PGNs are detected through peptidoglycan recognition proteins (PGRPs) that bind PGN and become activated (Masuzzo et al., 2022). In response to PGRP-mediated activation of Gr66a-expressing neurons there was a reduction in oviposition and feeding behavior in the adult fly. The mechanism involved in this response was shown to involve signaling through the IMmune-Deficiency (Imd) pathway, a canonical immune signaling pathway in flies (Masuzzo et al., 2022). Further confirming the ability of adult flies to sense bacteria, it was shown that in adult flies, Gr66a-expressing neurons display calcium signaling activity in response to the presence of PGN or *E. coli* in *in-vivo* experiments (Masuzzo et al., 2022). We sought to test whether fly larva could also detect bacteria by visualizing neuronal activity in larval taste neurons. We adapted a method previously used to analyze neuronal activity in adult flies following the addition of Gram-negative bacteria-derived PGNs (Masuzzo et al., 2022). This technique uses the reporter transgene *UAS-GCaMP6s* to monitor calcium signaling activity specifically in Gr66a-expressing neurons before and after the addition of different tastants. As a control, using bitter tasting quinine, we confirmed that Gr66a-expressing neurons show increased activity following bitter taste detection (Figure 1A-B; Figure 1 figure supplement 1A, and Video 1). In contrast, the addition of PGNs from Gram-negative bacteria did not result in an increase in calcium signaling activity in Gr66a-expressing neurons (Figure 1B). Importantly, experiments in which larvae were pre-incubated with PGNs and stimulated with quinine and PGNs revealed a dose-dependent inhibitory effect of PGNs on bitter sensing in Gr66a-expressing neurons (Figure 1C, and Figure 1 figure supplement 1B, and Video 2). The specificity of this effect was demonstrated by knocking down the PGN receptor PGRP-LC in *Gr66a-Gal4* neurons, which abolished the inhibitory effect of PGNs on the response to quinine (Figure 1D, and Figure 1 figure supplement 1C, and Video 3). Taken together, these results demonstrated that sensing of Gram-negative bacteria-derived PGNs by Gr66a-expressing neurons through the PGRP-LC receptor led to the inhibition of neural activity.

**Figure 1.**
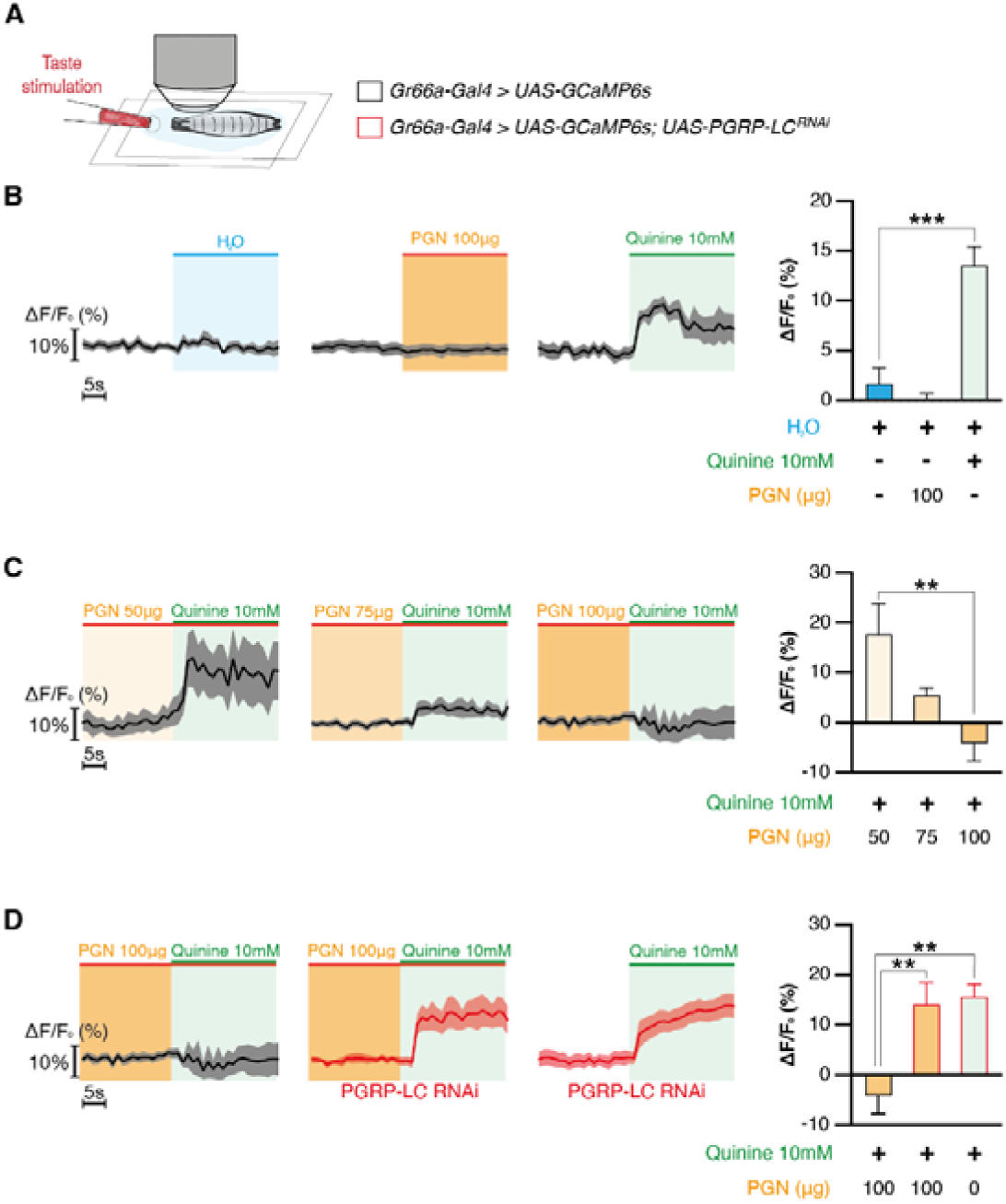
Peptidoglycan sensing inhibits Gr66a taste sensory neurons. (**A**), *In vivo* larvae receive taste stimulation directly under the microscope and Gr66a neural activity is monitored in control background or in PGRP-LC knocked down background. (**B**), Averaged curves of Gr66a neural activity in response to H_2_O (*n* = 8), peptidoglycan (*n* = 7), or quinine (*n* = 8) (left); and quantification of the responses (right). (**C**), Averaged curves of Gr66a neural activity in response to 10mM quinine preceded by peptidoglycan solutions at 50 μg/mL (*n* = 8), 75 mg/mL (*n* = 8), or 100 μg/mL (n = 8) (left); and quantification of the responses (right). (**D**), Averaged curves of Gr66a neural activity in response to 10mM quinine preceded by peptidoglycan solutions at 100 μg/mL in control larvae (*n* = 8), in larvae with PGRP-LC knocked down and in response to quinine alone (left); and quantification of the responses (right). Values represent mean ± SEM. Statistical tests: Kruskal-Wallis and Dunn’s post hoc; \*\**P* < 0.01; \*\*\**P* < 0.001.

### Taste modulates hematopoiesis

Having confirmed that larva can detect bacteria via taste we continued to ask whether taste, and more specifically, the activity of the bacteria sensing neurons, regulates larval hematopoiesis. To this end, we monitored blood cell differentiation and blood progenitor maintenance in third instar larval LGs whose most peripheral taste neurons were silenced by expression of the tetanus transgene under the control of the *Gr66a-Gal4* driver (*Gr66a-Gal4 > UAS-tnt*). By staining for the mature blood cell marker P1, we found that silencing bacterial taste sensory neurons resulted in an increase in the number of differentiated blood cells compared to controls (Figure 2A-B and Figure 2 figure supplement 1A-C) (Boll and Noll, 2002; LeDue et al., 2015). Interestingly, no modification in the number of crystal cells or lamellocytes was observed (Figure 2C, D; Figure 2 figure supplement 1E-G). Using *UAS-GFP* we confirmed that *Gr66a-Gal4* did not drive expression in any region of the LG (Figure 2 figure supplement 1H-H’’’) supporting the idea that it was indeed its ability to silence taste sensory neurons that led to increased blood cell differentiation. We did however see strong expression of *Gr66a-Gal4* in the peripheral taste organs (Figure 2 figure supplement 1). We confirmed that silencing peripheral aversive taste neurons induced increased plasmatocyte differentiation by silencing neurons from Gr33a-Gal4 which drives expression in an overlapping neuronal population in the peripheral taste sensory organs (Figure 2, figure supplement 2) (Tadres et al., 2025). Importantly, we confirmed that the increased plasmatocyte differentiation was unlikely to be the result of changes in metabolism by showing that the silencing of the Gr66a aversive taste neurons did not impact feeding (Figure 2 figure supplement 3). Taken together, these data strongly suggest that the impairment of peripheral aversive taste neurons leads to a specific increase in plasmatocyte differentiation in the LG without affecting feeding.

**Figure 2.**
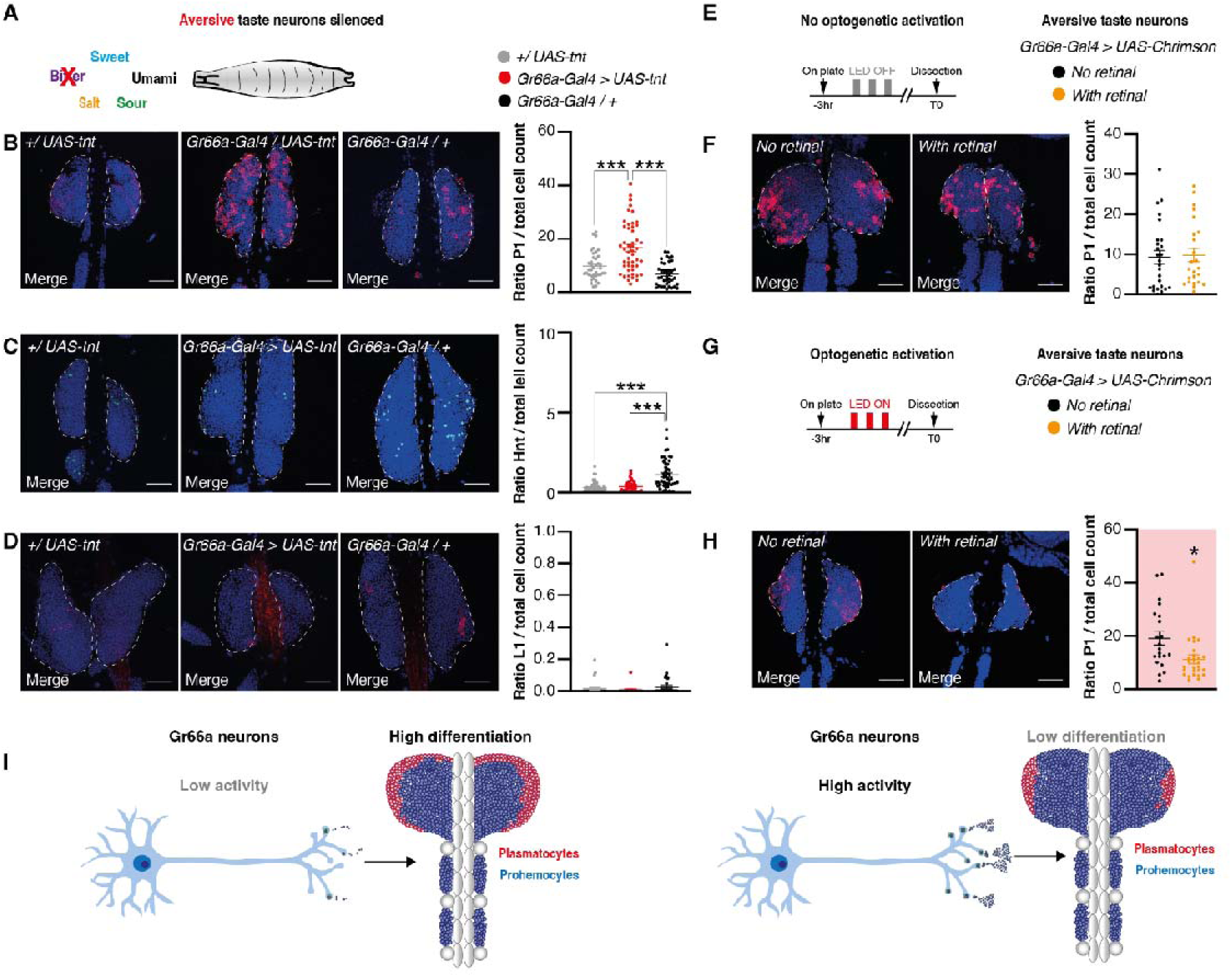
Aversive taste sensory neurons activity is oppositely correlated with plasmatocyte differentiation. (**A**), Schematic showing taste sensory modalities impaired in *Gr66a-Gal4>UAS-tnt* larvae. (**B**), Staining of lymph gland nuclei (Topro-blue) and differentiated plasmatocyte cells (left, P1-red) of *+/UAS-tnt* (*n* = 37), *Gr66a-Gal4>UAS-tnt* (*n* = 50), *Gr66a-Gal4/+* larvae (*n* = 39), and quantification of the ratio between the 2 (right). (**C**), Staining of lymph gland nuclei (Topro-blue) and differentiated crystal cells (left, Hnt-green) of *+/UAS-tnt* (*n* = 51), *Gr66a-Gal4>UAS-tnt* (*n* = 42), *Gr66a-Gal4/+* larvae (*n* = 49), and quantification of the ratio between the 2 (right). (**D**), Staining of lymph gland nuclei (Topro-blue) and differentiated lamellocyte cells (left, L1-red) of *+/UAS-tnt* (*n* = 26), *Gr66a-Gal4>UAS-tnt* (*n* = 16), *Gr66a-Gal4/+* larvae (*n* = 31), and quantification of the ratio between the 2 (right). (**E**), Schematic showing optogenetic protocol, the OFF light does not activate Gr66a neurons. (**F**), Staining of lymph gland nuclei (Topro-blue) and differentiated cells (left, P1-red) and quantification of the ratio between the 2 (right) in non-retinal fed larvae (*n* = 26), and retinal fed larvae (*n* = 24). (**G**), Schematic showing optogenetic protocol, the ON light activates Gr66a neurons in retinal flies. (**H**), Staining of lymph gland nuclei (Topro-blue) and differentiated cells (left, P1-red) and quantification of the ratio between the 2 (right)in non-retinal fed larvae (*n* = 18), and retinal fed larvae (*n* = 18). (**I**), When the activity of Gr66a-expressing is low, hematopoiesis is high (left). When the activity of Gr66a-expressing neurons is high, hematopoiesis is low (right). Scale bars are 50 μm. Values represent mean ± SEM. Statistical tests: one-way ANOVA and Tukey post hoc for A-D and *t*-test for E-H; \**P*< 0.05, \*\*\**P* < 0.001.

### Activation of Gr66a aversive taste neurons inhibits plasmatocyte differentiation

Since suppressing the activity of aversive taste neurons caused an increase in plasmatocyte differentiation in the LG, we asked whether the opposite treatment, consisting of activating these neurons using optogenetic tools, would modulate plasmatocyte differentiation. Specifically, we expressed the light-activated optogenetic transgene CsChrimson using *Gr66a-Gal4*. When larvae are fed with the cofactor all-trans retinal and exposed to red light CsChrimson causes neural activation (Klapoetke et al., 2014) (Figure 2E and 2F). We observed that activating Gr66a-expressing neurons (see materials and methods) promoted a reduction in plasmatocyte differentiation (Figure 2F and 2H, and Figure 2 figure supplement 4). These data demonstrate that the activity of Gr66a-expressing neurons is inversely correlated with plasmatocyte differentiation. When neuronal activity is low, plasmatocyte differentiation is elevated; conversely, increased activity is associated with reduced levels of plasmatocyte differentiation. (Figure 2I).

### Modulation of plasmatocyte differentiation by Gr66a-expressing neurons is mediated by PeptidoGlycan Receptor Proteins and a non-canonical Immune Deficiency pathway

Our data suggests that the activity of Gr66a-expressing neurons regulates larval hematopoiesis and that these neurons detect PGNs. Bacterial PGNs are typically detected through their interaction with members of the PeptidoGlycan family of transmembrane receptor proteins (Gottar et al., 2002; Lu et al., 2020). We asked whether PGRPs also control plasmatocyte differentiation. Intriguingly, previous studies have shown that PGRPs signal through the Imd pathway, which is required for antibacterial response of flies (Gottar et al., 2002; Lu et al., 2020; Yu et al., 2022). Although there are several PGRPs in flies that have distinct roles, the ability to signal through the Imd pathway is a shared feature. For example, PGRP-LC detects bacteria by binding to PGNs, and acts as a key regulator of the Imd pathway during the systemic immune response in the fat body (Yu et al., 2022). Moreover, PGRP-LA and PGRP-LF, neither of which is known to directly bind PGNs, act as positive and negative regulator of the Imd pathway, respectively (Basbous et al., 2011; Gendrin et al., 2013; Maillet et al., 2008; Yu et al., 2022). Binding of PGNs to the PGRP-LC receptor regulates several protein components of the Imd signaling pathway including Imd, Fadd, Dredd, and controls the nuclear localization of the NF-kB transcription factor Relish (Yu et al., 2022). To determine whether Gr66a-expressing neurons sense PGNs through PGRPs and signal through the Imd pathway, we used RNAi-mediated knockdown for various PGRPs and Imd components (Figure. 3A). We found that targeted knock-down of the PGRP-LC receptor in Gr66a neurons resulted in a reduced plasmatocyte differentiation (Figure 3, and Figure 3 figure supplement 1A-C). Additionally, knockdown of the positive Imd pathway regulator, PGRP-LA, also led to reduced plasmatocyte differentiation. In contrast, knockdown of the negative Imd pathway regulator, PGRP-LF, increased plasmatocyte differentiation (Figure 3, and Figure 3 figure supplement 1A-C) as well as amplified the inhibitory effect of PGN to the response to quinine (Figure 1 figure supplement 1D). In line with these observations, knockdown of downstream components of the Imd pathway such as Dredd and Fadd also decreased plasmatocyte differentiation (Figure 3 figure supplement 1D-J). Interestingly, although Relish is canonically regarded as the rate-limiting effector of the Imd pathway, its knockdown using two different RNAi lines did not alter plasmatocyte differentiation. This suggests that Imd signaling in Gr66a neurons may occur via a non-canonical pathway (Figure 3, and Figure 3 figure supplement 1A-C).

**Figure 3.**
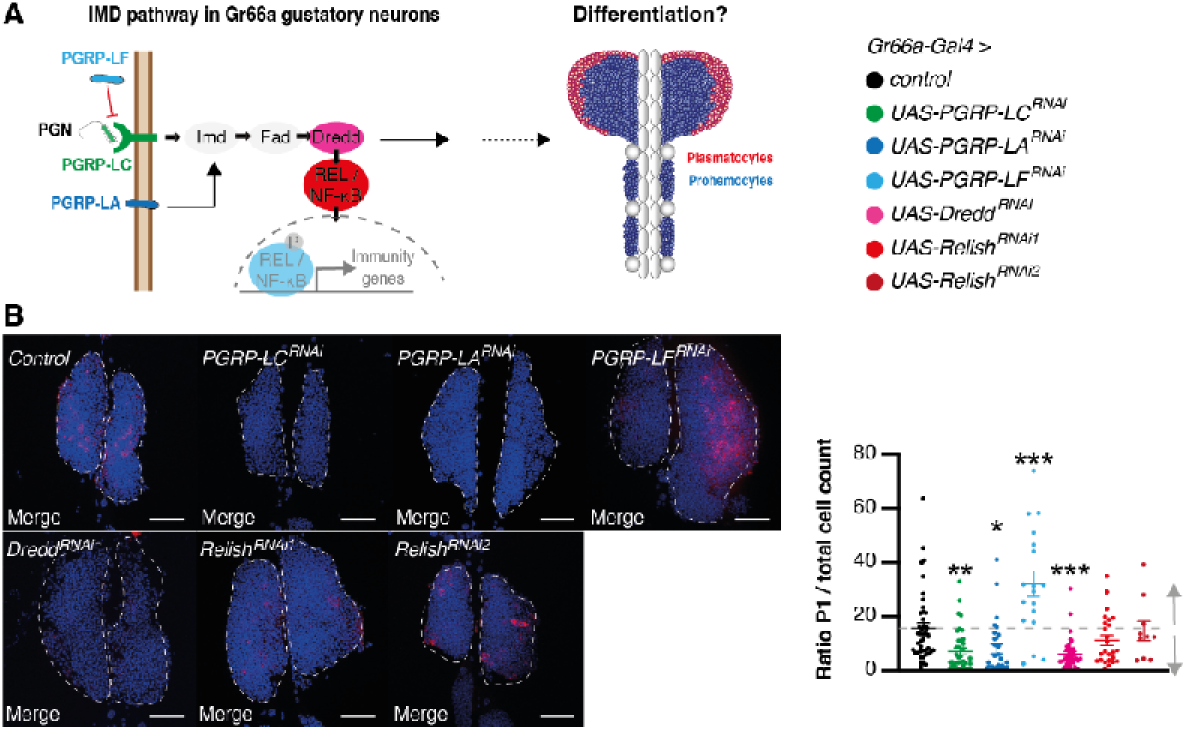
Knocking down PGRPs and Imd pathway in Gr66a taste sensory neurons inhibits hematopoiesis. (**A**), Schematic showing how peptidoglycan detection by the Imd pathway in Gr66a taste sensory neurons could regulate hematopoiesis. (**B**), Staining of lymph gland nuclei (Topro-blue) and differentiated cells (left, P1-red) of *Gr66a-Gal4 > Imd ^RNAi^* larvae (Control, *n* = 42; PGRP-LC, *n* = 38; PGRP-LA, *n* = 30; PGRP-LF, *n* = 18; Dredd, *n* = 39; Relish, *n* = 25, and 10), and quantification of the ratio between the 2 (right). Scale bars are 50 μm. Values represent mean ± SEM. Statistical tests: one-way ANOVA and Tukey post hoc; \**P* < 0.05; \*\**P* < 0.01; \*\*\**P* < 0.001.

Since, Relish knockdown did not alter plasmatocyte differentiation, despite its canonical role as key Imd pathway effector, we asked whether other immune signalling pathways might be involved in signalling in Gr66a neurons downstream of PGN detection. Knocking-down Spätzle and cactus, two essential components of the Toll pathway, did not impact plasmatocyte differentiation (Figure 3 figure supplement 2). However, since previous work demonstrated cross-talk between the Imd and JNK pathways (Maillet et al., 2008) we also tested key JNK components. Interestingly, knockdown of two JNK pathway kinases (basket and Hep) reduced plasmatocyte differentiation (Figure 3 figure supplement 2). Finally, we demonstrated that modulation of the Imd pathway in Gr66a neurons did not impact larval feeding (Figure 3 figure supplement 3), demonstrating that the plasmatocyte differentiation modulatory effect is unlikely to be a result of changes in nutrition due to altered food consumption but rather from modification of Gr66a signaling.

Our data suggest that aversive taste neurons expressing the Gr66a taste receptor regulate plasmatocyte differentiation through PGRP-LC-mediated signalling. However, this does not exclude the possibility that other sensory receptors in these neurons might also contribute to this process. Gr66a-expressing neurons are known to detect both bitter compounds via the gustatory receptor Gr66a (Apostolopoulou et al., 2015; Marella et al., 2006; Wang et al., 2004) and, lipopolysaccharides (LPS), bacterial endotoxins that are known to activate innate immunity via the chemosensory cation channel dTRPA1 (Chen and Dahanukar, 2020; Soldano et al., 2016). To test whether bitter and LPS sensing modulate plasmatocyte differentiation, we knocked down both the Gr66a and dTRPA1 receptors using RNAi. Neither manipulation altered plasmatocyte differentiation relative to controls (Figure 3 figure supplement 4). Taken together, these findings suggest that plasmatocyte differentiation is controlled by a non-canonical Imd-dependent mechanism in Gr66a-expressing taste neurons, and that occurs independently of standard bitter or LPS sensing.

### Detection of Gram-negative bacteria through PGRP-LC regulates blood cell differentiation

Our results support a model in which Gr66a-expressing neurons detect PGNs, activating the Imd pathway and modulating plasmatocyte differentiation. We asked if this mechanism operates under physiological conditions of bacterial exposure. Specifically, we investigated whether Gr66a-expressing neurons detect Gram-negative bacteria via PGN and, if so, this would promote immune cell production *in-vivo* in the adult fly. To this end, we raised larvae in an antibiotic enriched (ATB) media to minimize microbial exposure (see Materials and Methods), and subsequently exposed them to *Escherichia coli (E.coli),* a Gram-negative bacterium (Figure 4A). Consistent with previous observations, we found that exposure to *E. coli* was sufficient to induce an increase in plasmatocyte differentiation (Figure 4B, and Figure 4 figure supplement 1).

**Figure 4.**
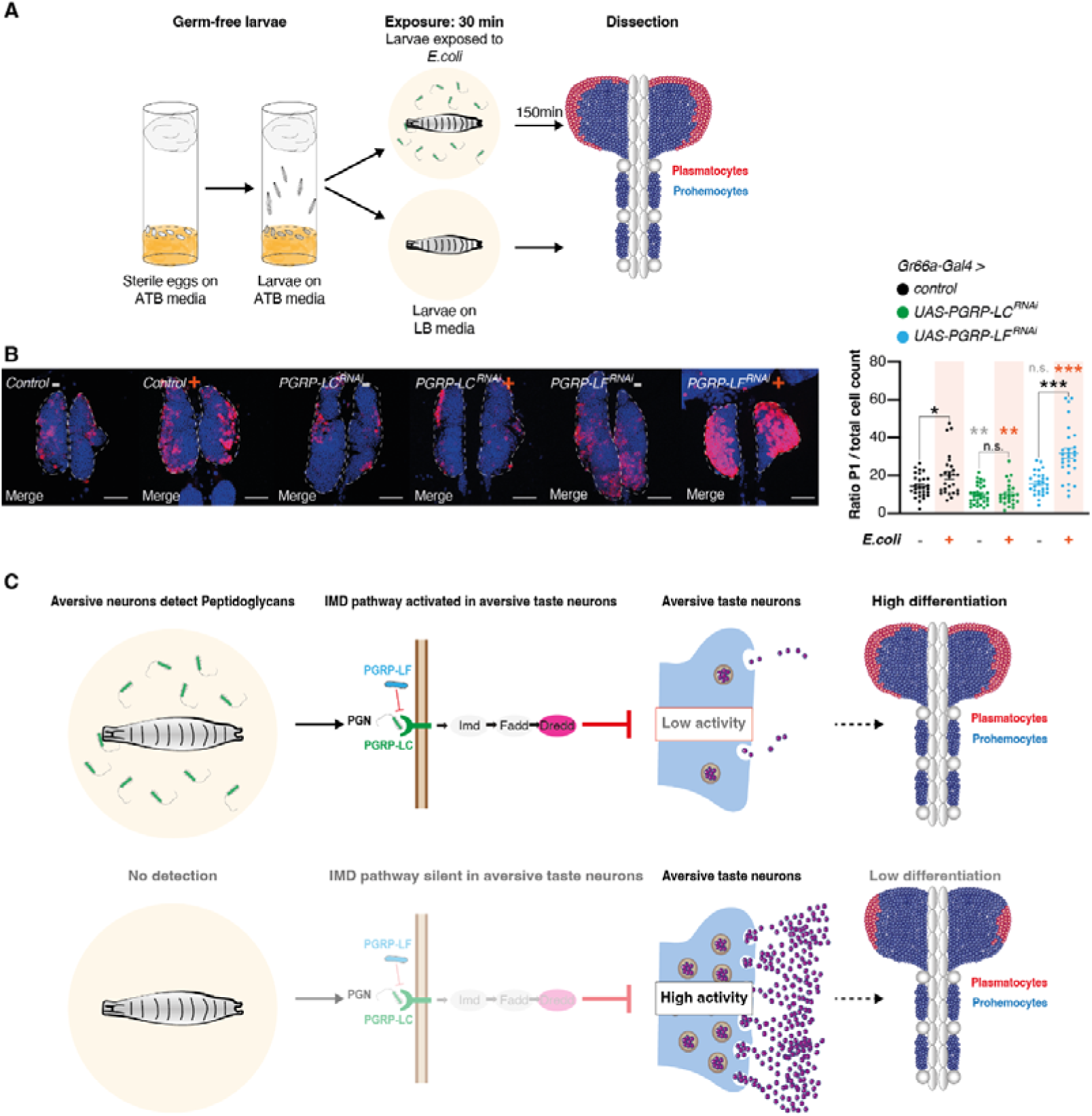
PGRP-LC in Gr66a gustatory neurons mediates increased hematopoiesis after tasting bacteria. (**A**), Schematic representation of the protocol for the exposure of germ-free larvae with *E. coli*. After oviposition the embryos grow under sterile conditions in vials containing ATB media. Then larvae are transferred to LB plates containing either *E. coli* or nothing for 30 minutes and placed back to LB media for 150 minutes before dissection. (**B**), Staining of lymph gland nuclei (Topro-blue) and differentiated cells (left, P1-red) of *Gr66a-Gal4 > Imd ^RNAi^* flies without exposition to *E. coli* (Control, *n* = 28; PGRP-LC, *n* = 32; PGRP-LF, *n* = 28) and with exposition to *E. coli* (Control, *n* = 27; PGRP-LC, *n*= 24; PGRP-LF, *n* = 28), and quantification of the ratio between the 2 (right). (**C**), In the presence of gram-negative bacteria, larvae detect peptidoglycans through the PGRP-LC/Imd signaling pathway from Gr66a gustatory neurons, which inhibits their activity, thus leading to an increased hematopoiesis. In the absence of Gram-bacteria, the PGRP-LC/Imd signaling pathway from Gr66a gustatory neurons is not activated, which does a release of their inhibition, thus leading to a decreased hematopoiesis. Scale bars are 50 μm. Values represent mean ± SEM. Statistical tests: one-way ANOVA, Tukey post hoc, and unpaired *t*-test for comparison of the same genotype between exposed and non-exposed group; \**P* < 0.05; \*\**P* < 0.01; \*\*\**P* < 0.001.

We next asked whether this response required signalling via the PGN receptor PGRP-LC in Gr66a-expressing neurons. Knockdown of PGRP-LC reduced plasmatocyte differentiation both in the presence and absence of bacteria, suggesting this receptor has a basal level of activity even when bacteria are not present (Figure 4B, and Figure 4 figure supplement 1). Conversely, knockdown of the inhibitory PGRP-LF, which is expected to enhance PGRP-LC activity, increased plasmatocyte differentiation, but only in the presence of bacteria (Figure 4B, and Figure 4 figure supplement 1). This suggests that PGRP-LF acted as an inhibitor of PGRP-LC but only in the presence of bacteria. Taken together, these results show that, in Gr66a-expressing neurons, modulation of PGRP-LC signalling controlled plasmatocyte differentiation in a bacteria-dependent manner (Figure 4C, and Figure 4 figure supplement 1).

### Sensory detection of Gram-negative bacteria primes the immune system

Our findings were consistent with a model in which sensory detection of Gram-negative bacteria promotes hematopoiesis, specifically enhancing plasmatocyte production. Plasmatocytes play a critical role in bacterial defense, and their absence leads to increased susceptibility to several microorganisms and reduced survival following infection (Charroux et al., 2009). We asked whether the increase in plasmatocyte differentiation triggered by sensory detection of *E. coli* was sufficient to confer improved resistance to bacterial infection. To test this, we reared larvae in germ-free condition (see Materials and Methods) and exposed them briefly to *E. coli*. After pre-exposure, we returned the larvae to antibiotic-containing vials to continue development. Adults that emerged from these treatments were infected with *Pseudomonas entomophila* by puncture, and their survival was monitored (Figure 5A).

**Figure 5.**
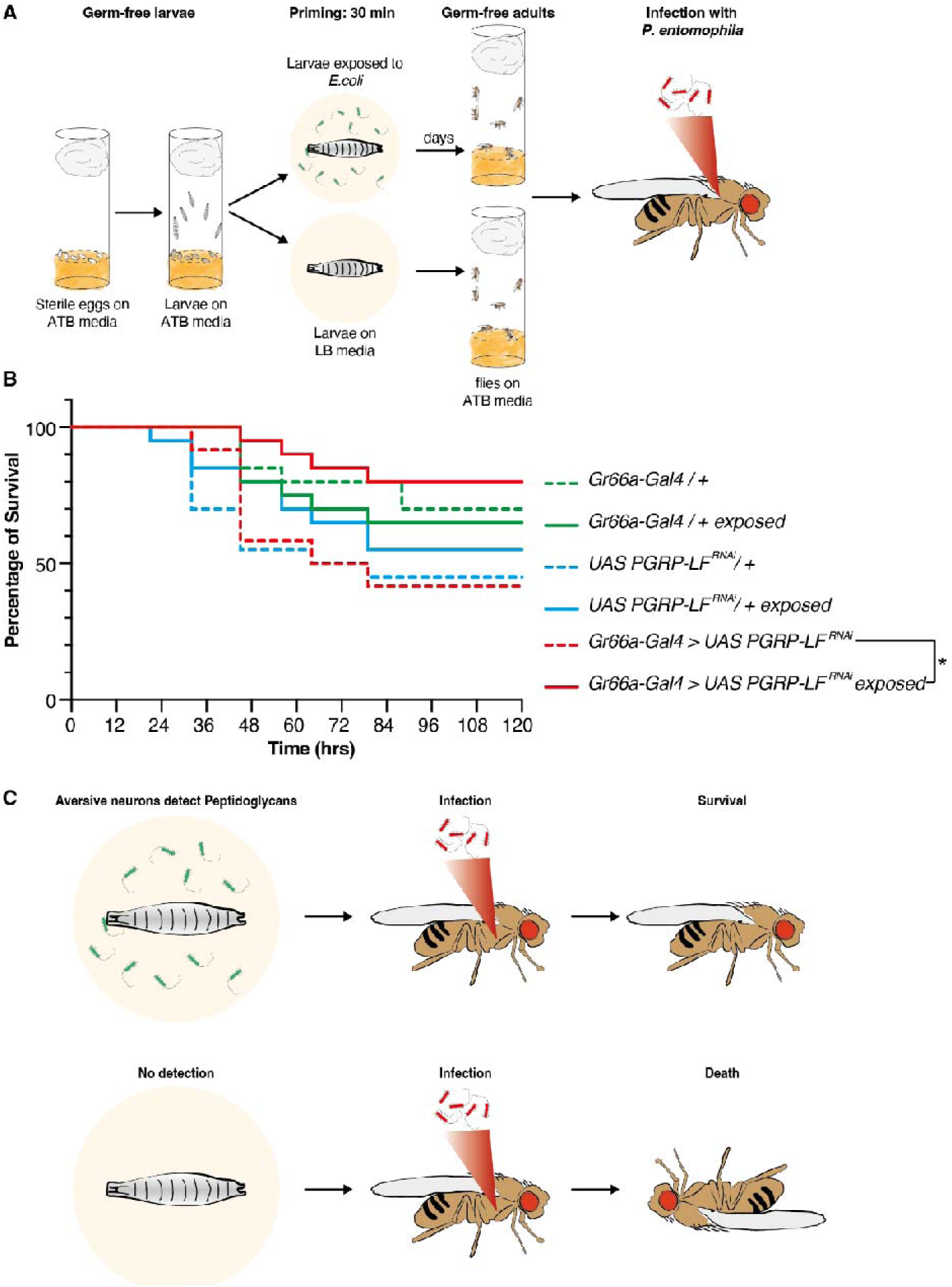
Overactivation of Imd pathway in Gr66a gustatory neurons primes the cellular immune pathway. (**A**), Schematic representation of the protocol for the exposition of germ-free larvae with *E. coli* followed by infection with *P. entomophila* in the adults. After oviposition the embryos grow under sterile conditions into a vial containing ATB media. Then larvae are transferred to LB plates containing either *E. coli* or nothing for 30 minutes and placed back to LB media for 150 minutes. Then the larvae are placed on ATB media until adulthood. Adult flies are then infected by *P.entomophila* and their survival is monitored. (**B**), Survival curves following infection with *P.entomophila* of flies previously exposed to *E.coli* (solid lines, *Gr66a-Gal4 > +, n* = 20; *+ / UAS-PGRP-LF^RNAi^*, *n* = 20; *Gr66a-Gal4 > UAS-PGRP-LF^RNAi^*, *n* = 19) or to nothing (dashed lines, *Gr66a-Gal4 > +*, *n* = 20; + */ UAS-PGRP-LF^RNAi^*, *n* = 20; *Gr66a-Gal4 > UAS-PGRP-LF^RNAi^*, *n* = 12). (**C**), In the presence of gram-negative bacteria, the activation of the Imd signaling pathway from Gr66a gustatory neurons, promotes higher resistance to infection in adult flies. Values represent mean. Statistical tests: one-sided log-rank test; \*\**P* < 0.01.

To avoid inducing a large-scale systemic immune response, the pre-exposure period was deliberately kept short resulting in only a modest (a 6%) increase in plasmatocyte numbers (Figure 5). Consequently, we found that infected adults pre-exposed to *E. coli* as larva did not fare any better than infected adults that were not pre-exposed (Fig. 5B, and Figure 5 figure supplement 1). Similarly, the mild reduction in plasmatocyte numbers caused by PGRC-LC knockdown in Gr66a-expressing neurons in larvae (∼15%, Figure 4) did not alter adult survival (Fig. 5B, and Figure 5 figure supplement 1).

By contrast, knockdown of the inhibitory receptor, PGRP-LF in Gr66a-expressing neurons, produced a substantially larger increase in plasmatocyte numbers (up to 31%) following pre-exposure to *E. coli*. Importantly, these larvae exhibited improved adult survival following septic infection (Fig. 5B, and Figure 5 figure supplement 1). These findings suggest that larval exposure to bacteria can confer lasting resistance to infection into adulthood, but only when it induces a sufficiently large, taste-mediated increase in plasmatocyte production. (Fig. 5C).

## DISCUSSION

Animals surveil their environment in search of cues that inform decisions that control feeding and warn of potential threats. Here we ask whether such environmental cues can act to proactively regulate immune defenses prior to a pathogenic infection. Specifically, we show that taste perception is able to directly regulate hematopoiesis by controlling plasmatocyte differentiation in Drosophila larvae. We demonstrate that Gr66a-expressing sensory neurons, canonically associated with aversive-bitter taste detection, also function as bacterial sensors by detecting peptidoglycans (PGNs) through peptidoglycan recognition protein LC (PGRP-LC). Rather than simply conveying taste information, these neurons can act as immunomodulatory sensors that engage a non-canonical Imd pathway to regulate plasmatocyte differentiation. Activation of these neurons suppresses hematopoiesis, while their inhibition, either by PGN stimulation or genetic manipulation, promotes plasmatocyte production. Importantly, short-term bacterial exposure during larval stages leads to increased immune cell production and confers enhanced resistance to infection in adulthood via the PGRP pathway in Gr66a-expressing sensory neurons. Together our findings reveal that taste is not only a sensory modality to evaluate food quality, but an anticipatory immune sensing system that prepares the organism for infection risk.

Taste receptor-mediated pathogen detection is well documented in mammals. Specialized, epithelial immune-sensing cells, such as solitary chemosensory cells in the mucosa, brush cells in the respiratory tract and tuft cells in the gut, express taste receptors and detect bacterial components (Xi et al., 2022). Upon detecting a pathogen, these cells locally release antimicrobial peptides and cytokines, engaging humoral immune defenses (Xi et al., 2022). Notably, these examples of taste receptor-mediated responses are typically localized and have not been shown to involve information transfer through neural circuits (Xi et al., 2022). In contrast, Gr66a-expressing neurons in Drosophila are bona fide neural sensors, they are located in peripheral taste organs whose organization is similar to that of mammalian taste buds (Yarmolinsky et al., 2009), send axonal projections transmitting sensory information to higher brain structures (Engert et al., 2022; Kim et al., 2017; Talay et al., 2017; Weber et al., 2023), and act at a distance rather than through local immune secretion. This raises the possibility that other sensory modalities may regulate hematopoiesis through additional neural pathways. Although the specific mechanism by which sensory information is being relayed from the brain to the lymph gland (LG) remains unresolved, several neuroimmune signaling pathways are known to regulate hematopoiesis in *Drosophila* (Goyal et al., 2022; Rodrigues et al., 2021; Shim et al., 2013; Yu et al., 2022). These pathways, that include brain-derived *Drosophila* insulin-like peptides (DILPS), GABAergic signaling, and serotonin signaling via the 5-HT1B receptor (Bland, 2023; Goyal et al., 2022; Shim et al., 2013), provide plausible mechanisms through which sensory neural activity could influence immune cell development. Identifying which of these signals act downstream of taste detection will be a key focus for future work.

Previous studies have demonstrated some sensorial cues can regulate hematopoiesis, and that each sensory modality produces a distinct immune outcome. For example, olfactory cues associated with food quality promote maintenance of hematopoietic progenitors through neuronal signaling (Goyal et al., 2022; Madhwal et al., 2020; Shim et al., 2013), while odours associated to parasitic pathogens detected by the olfactory system induce lamellocyte differentiation in anticipation of wasp infestation (Madhwal et al., 2020; Shim et al., 2013). These studies established that sensory systems can shape immune cell fate depending on the nature of the environmental threat. Our work extends this principle to the gustatory system by showing that detection of Gram-negative bacteria through taste promotes plasmatocyte differentiation. This suggests that individual sensory modalities may selectively instruct distinct hematopoietic fates, enabling animals to tailor their immune responses to the anticipated type of pathogen. Given that *Drosophila* encounter diverse microbial threats such as Gram-positive bacteria and fungi, we hypothesize that sensory modalities may help mount a unique anticipatory immune response to specific immune challenges.

To investigate how sensory input is functionally translated into hematopoietic output, we examined how the activity of Gr66a-expressing neurons relates to plasmatocyte differentiation. Our results reveal that the level of activity in Gr66a-expressing neurons influences hematopoiesis: reducing neuronal activity promotes plasmatocyte differentiation whereas increasing their activity decreases it. Gr66a-expressing neurons are polymodal sensory neurons that express a variety of receptors responsive to a broad range of chemical cues, including bitter compounds (Gr66a, Gr33a), high salt (Sano, ppk19), bacterial lipopolysaccharides (TrPA1) (Alves et al., 2014; Kwon et al., 2011; Soldano et al., 2016). Or even electric fields (Tadres et al., 2025). Consequently, knocking down the Gr66a receptor disrupts bitter sensing specifically but leaves other sensory pathways active, whereas silencing Gr66a neurons altogether eliminates input from all sensory modalities and lowers overall neuronal activity. These observations suggest two non-mutually exclusive models by which sensory neurons could regulate hematopoiesis: (1) plasmatocyte differentiation may be induced when the overall activity of Gr66a-expressing neuron falls below a certain threshold, or (2) differentiation may be triggered by the inhibition of specific sensory pathways within these neurons. Our data support the second model, as exposure to bacterial peptidoglycan (PGN) selectively inhibits Gr66a neuronal activity via PGRP-LC and promotes plasmatocyte differentiation.

Because they regulate both taste perception and hematopoiesis, we considered whether microbial detection by Gr66a neurons might also influence feeding behaviour. In larvae, silencing Gr66a disrupts bitter sensing, but not sweet preference, and does not impair feeding on standard or yeast-based food (Apostolopoulou et al., 2014; Maier et al., 2021). Consistent with this we found that PGN exposure inhibits Gr66a neuronal activity, suggesting that larvae are less likely to reject bacterially contaminated food, even if it contains bitter compounds. Our imaging data shows that the detection of bacteria-derived PGN inhibits the activity of bitter-sensing neurons, meaning that larvae should be less likely to reject a bitter-tasting food if it is contaminated with bacteria. In contrast, in adult flies, PGN detection has been shown to activate bitter-sensing neurons and promote food avoidance (Masuzzo et al., 2022). Moreover, the aversive response to bacterial PGNs in adults requires prior exposure, or “priming” during larval development, and enhanced calcium activity in specific neurons following PGN exposure also depends on Gr66a signaling (Montanari et al., 2024). Thus, flies raised under conventional conditions display PGN-mediated food avoidance, whereas axenic flies do not. These observations raise an evolutionary question: why would larvae suppress bitter rejection when encountering bacterially contaminated food? One possibility is that early-life exposure to bacterial cues may be beneficial by promoting hematopoiesis and establishment of the microbiota, thereby strengthening immune competence later in life. Indeed, our findings support this idea, as larvae that experience bacteria sensing exhibited enhanced plasmatocyte production and increased resistance to infection in adulthood.

Immunological memory has traditionally been attributed to the adaptive immune system (Medzhitov and Janeway, 2000), yet accumulating evidence suggests that innate immunity can also exhibit forms of memory, often referred to as “immune priming” or “trained immunity” (Arch et al., 2022; Aymeric et al., 2010). In our study, we show that sensory detection of bacterial peptidoglycans (PGNs) via the Imd pathway in Gr66a-expressing neurons promotes plasmatocyte differentiation during larval stages and that larvae with a sufficiently elevated number of plasmatocytes display greater resistance to infection as adults. This demonstrates that a prior exposure to bacteria, even in the absence of infection, can confer protection in the future, resembling immune priming (Arch et al., 2022). Previous studies have shown induced immune priming in *Drosophila* by infecting adult flies through puncture or ingestion, triggering both cellular and humoral responses. By contrast, our findings reveal that priming can occur *without* systemic infection and can be initiated solely by sensory perception of microbial cues. We propose that this form of anticipatory immune activation represents a distinct mechanism, which we term Taste Primed Immunity (TPI). TPI differs from previously described forms of immune priming in two key ways. First, it is initiated during larval development rather than in adults. Second, it acts by modulating the cellular branch of immunity, specifically by promoting plasmatocyte differentiation. Since up to 90% of circulating plasmatocytes in adults are derived from larval hematopoiesis (Bosch et al., 2019; Koranteng et al., 2022), taste-mediated enhancement of larval hematopoiesis has lasting consequences for immune competence in adulthood.

In summary, our work reveals that taste is not merely a chemosensory system but functions as an anticipatory immune sensor that detects bacterial signatures and proactively modulates hematopoiesis. Through a non-canonical Imd pathway in Gr66a-expressing neurons, larvae adjust immune cell production in response to sensorial prediction of infection risk, shaping both immediate and long-term immune resilience. We propose that this mechanism represents a form of sensory-driven predictive immunity, which we term Taste-Primed Immunity (TPI).

## MATERIALS AND METHODS

### Drosophila melanogaster

Fly stocks were raised on standard food at 25C and 70% relative humidity under a 12-hour light/12-hour dark cycle. For neuronal silencing experiments, we used *UAS-tnt*. For neuronal activation experiments, we used *20XUAS-IVS-CsChrimson.mVenus* (Bloomington #55135). Bitter taste sensory neuron expression was driven by *Gr66a-Gal4* (*32*) and *Gr33a-Gal4* (Bloomington#31425). For staining experiments, we used *40XUAS-IVS-mCD8::GFP* (Bloomington#32195), For imaging experiments, we used the *20XUAS-IVS-GCaMP6s* (Bloomington #42746). For RNAi experiments, we used RNAi against Gr66a (Bloomington #31284), TrpA1 (Bloomington #66905), PGRP-LC (VDRC #101636KK), PGRP-LA (VDRC #102277KK), PGRP-LF (VDRC #108313KK), Dredd (VDRC #104726KK), Relish (VDRC #108469KK for RNAi1 and 49413GD for RNAi2), PGRP-LB (Bloomington #67236), PGRP-LD (Bloomington #34892), PGRP-LE (Bloomington #60038), Fadd (Bloomington #34813), PGRP-SB2 (Bloomington #56982), PGRP-SB1 (Bloomington #56983), PGRP-SC1b (Bloomington #67249), PGRP-SC2 (Bloomington #56315), bsk (Bloomington #35594), hep (Bloomington #28710), spz (Bloomington #28538), cactus (Bloomington #37484). Controls for RNAi experiments (Bloomington #35784; VDRC #60000TK).

### Embryo sterilization

Embryo sterilization was performed as previously (Montanari et al., 2024). Three petri dishes are filled with 2.6% bleach, 70% ethanol (Ethanol 96° RPE Carlo Erba Ref 414638) and autoclaved purified distilled water respectively. The embryos are collected by filling the tube in which oviposition occurred with PBS and using a small brush to gently detach them from the flies’ food. A 40μm cell strainer is used to collect the embryos. The cell strainer with the embryos is then dipped into: bleach 2.6% for 5 minutes, ethanol 70% for 1 minute, purified water for 1 minute, ethanol 70% for 1 minute, purified water for 1 minute. The brush used to collect the embryos is sterilized in 2.6% bleach for ten minutes, rinsed and then used to transfer the sterile embryos on antibiotic enriched media.

### Larval exposure protocol

Early third instar larvae from sterile embryos were placed in petri-dish containing LB media (non-exposed group) or in petri-dish containing LB media contaminated with *E. coli* for thirty minutes and then placed on petri-dish containing LB media for two hours and a half until dissection (exposed group).

### Larval feeding protocol

We used a previously published protocol (Bhatt and Neckameyer, 2013). Third instar larvae were collected and transferred to the center of either an agar-filled plate overlaid with 5 ml of 2% activated baker’s yeast solution, or to the center of a standard-food plate. The rate of mouth hook contractions directly correlates with the amount of food ingested. The larvae were acclimated for 30 sec. then observation and recording of the number of mouth hook contractions for a period of 1 min were performed. *n* = 30 for each genotype.

### Dye ingestion protocol

Food-intake assays were performed using adapted protocol (Maier et al., 2021). L3 Larvae were starved for 4 hours at 25°C on 1% agarose petri dishes as a water source to keep the inside of the container humidified. Behavioural tests used active yeast as food source. A blue food dye (Erioglaucine Disodium Salt, Sigma 861146) was added to the yeast at a 2:1 ratio. Starved larvae were carefully transferred into petri dishes with dyed yeast to start the feeding process for one hour. Larvae were homogenized in 200□μl PBS buffer in 1.5□mL Eppendorf tubes with a tungsten bead, and grinded 30s at 30 Hz (3 times). Products were centrifuged twice (10 000□r.p.m.) for 10□min to clear the debris. After centrifugation, the absorbance of the supernatant was measured at 630□nm (A630). The background absorbance for supernatants from flies fed with regular food was subtracted from the absorbance of the supernatant from blue food-fed flies. The net absorbance reflected the amount of food ingested. A *n* of 1 correspond to 10 larvae.

### Dissections and Immunohistochemistry

Third instar larvae were collected and prepared for lymph gland dissection in cold 1% phosphate buffered saline (PBS) and fixed in 4% paraformaldehyde (PFA) for 15 minutes, then washed two times for 5 minutes with 0.1% PBST (PBS with 0.1% Triton-X [ThermoFisher Scientific BP151100]). The blocking step was done for 15 minutes using 16% Normal Goat Serum (NGS). Samples were then incubated overnight at 4°C in a 1:100 dilution of anti-P1 (NimC P1 a+b) in 0.1% PBST primary antibody (gift from Dr. Istvan Ando, Hungarian Academy of Sciences, Hungary). After overnight incubation, the samples were washed twice for 5 minutes using 0.1% PBST and then blocked for 15 minutes with NGS. Secondary antibody incubation of the samples was done for 2 hours at room temperature using a solution of donkey anti-mouse cy3 (1:400, Code: 715-165-150, RRID: AB_2340813, Jackson Immunoresearchlaboratories) and TO-PRO diluted in 0.1% PBST, followed by a set of three, 5-minute washes with 0.1% PBST. LG samples were mounted in glass bottom mounting dishes using VECTASHIELD Antifade Mounting Medium with TO-PRO (H-1200, Vector Laboratories, RRID:AB2336790). For crystal cell and lamellocyte stainings, samples were incubated overnight at 4°C in a 1:50 primary antibody dilution of anti-Hnt (DSHB; 1G9-s) or a 1:25 dilution of anti-L1 (gift from Dr. Istvan Ando, Hungarian Academy of Sciences, Hungary) respectively, with 0.1 % PBST. After incubation, the samples were washed twice for 5 minutes in 0.1% PBST and blocked for 15 min in Neat Goat Serum (NGS). Secondary antibody incubation was done for 2 hours at room temperature with a solution of donkey anti-mouse cy3 (1:400, Code: 715-165-150, RRID: AB_2340813, Jackson Immunoresearchlaboratories) and TO-PRO diluted in 0.1% PBST, followed by three, 5-minute washes with 0.1% PBST."

### Survival experiments

LB agar plates were seeded with 100 µL of an overnight *E. coli* culture and incubated overnight at 37°C. Wandering third instar larvae were placed for 30 minutes on LB agar plates coated or not with *E. coli* and then transferred onto LB only plates for 2h30. By batch of 20, larvae were transferred into vials containing fly food supplemented with antibiotics placed in 25°C incubators until fly hatching. Hatched 5-to-7-day old females were puncted with a tungsten needle dipped into an OD10 solution of *P. entomophila*.

### Imaging

For Lymph gland imaging, after overnight incubation, mounted LG samples were imaged using an Olympus FV1000 inverted confocal microscope with a numerical aperture 1.30 UPLFLN with the 40x oil immersion lens. Laser lengths used for detection of staining were 559 nm for Cy3 and 635 nm for Cy5. LG images were taken along the z-axis, 3nm apart.

For neuron imaging, larvae were partly dissected in PBS, fixed in 4% paraformaldehyde for 45 min at room temperature (RT). Anterior part of the larvae and guts were then isolated by precise dissection. They were mounted in VECTASHIELD® mounting medium (H-1000, Vector Laboratories). Fluorescence was observed with a Leica TCS SP8 confocal microscope.

### *In vivo* calcium imaging

*In vivo* larval calcium imaging experiments were carried out on wandering third instar larvae. Larvae were immobilized in a small drop of distilled water placed between two plastic coverslips (22 mm x 22 mm, Agar Scientific). The two coverslips were held together to prevent movement of the anterior part of the larvae, using alligator clips attached to a support. Stimulation was performed manually using a pipette with gel loading tip by applying water, PGN (50µg to 100µg/mL) or quinine (10 mM) diluted in water through a hole made in the upper coverslip allowing the solutions to come into contact with the Terminal Organ (TO) of the larvae. In PGN preincubation experiments, larvae were preincubated with the indicated dilution of PGN for one or two minutes and a solution containing the appropriate dilution of PGN and 10mM quinine was applied. Gr66a neurons were recorded in the TO of heterozygous larvae (*GR66a-Gal4>UAS-GCaMP6s*) with a 10x dry objective. GCaMP6s was excited using a Lumencor diode light source at 482 nm ± 25. Emitted light was collected through a 505-530 nm band-pass filter. Images were collected every 500ms using a Hamamatsu/HPF-ORCA Flash 4.0 camera and processed using Leica MM AF 2.2.9. Each experiment consisted of a recording of 70-100 images before stimulation and 160 images after stimulation. Data were analyzed as previously described (Masuzzo et al., 2022; Montanari et al., 2024) by using FIJI (https://fiji.sc/). For larvae fluorescence quantifications, a background fluorescence variation was calculated and subtracted to the fluorescence variation signal. For each image, ROIs were manually drawn on the left and right side and the signal intensity was quantified. These data were then manually inspected to exclude images with clear signs of drift and to select the side with the least variation in response. The initial intensity F_0_ was calculated over 10 frames, 30 frames before the stimulus, and ΔF/F_0_ was expressed as a percentage. Peak ΔF/F_0_ was calculated as the average ΔF/F_0_ over 4 frames around the peak minus average ΔF/F_0_ over 4 frames immediately before the peak. Statistical differences in peak ΔF/F_0_ between species were assessed using the Mann–Whitney U test. For calcium imaging experiments, Peptidoglycan (PGN) was purchased from InvivoGen (PGN-EK, # tlfl-pgnek) and quinine from Sigma-Aldrich (# Q1125).

### Optogenetics

The optogenetics experiments set up in this study were designed based on a previously published protocol by Honjo (Honjo et al., 2012). Genetic crosses were set up to yield *Gr66a-Gal4 / UAS-Chrimson* and *PLB-Gal4 / UAS-Chrimson* lines. Larvae were kept in vials and growing in the dark as Chrimson is a photosensitive channel that might be deactivated due to light exposure. For both genotypes, two groups of larvae were made respectively. One group was fed fly food that contained 2.5 µL of 100 µM all-trans-retinal (Ret+), a cofactor of channelrhodopsin. The second group of larvae were fed food containing 2.5 µL of ethanol (Ret-). Ret+ and Ret-groups of larvae were subjected to one of two conditions, they were either exposed to three cycles of 15 minutes of red LED light (671 nm) with 15 minutes in between cycles where they were kept in the dark, or they were not exposed to LED light at all. To allow time for hemocyte differentiation LGs were dissected two hours after the last cycle.

### Glucose and Glycogen Assay

For glucose and Glycogen assay, we adapted a previously published protocol (Wat et al., 2020). 10 larvae L3 were collected in a pre-weighed 1.5 mL tubes and stored at −80°C until processed. Before starting the assay, we weighed the tubes containing the larvae as they come out of the freezer. Larvae were then homogenized using 3 x 2mm Zirconium oxide beads (Next Advance – ZROB20) in 200 μl PBS 1X with a tissue lyser (Qiagen) (3X1 min at 30hz). There were centrifuged (Eppendorf centrifuge 5430R) a first time for 10 min at 10000 rpm at 4°C. For each sample, 50 μL of homogenate were transferred to a 1.5mL eppendorf and heat treated at 70°C for 10 min. They were again centrifuged for 7 min at 10000 rpm at 4°C to precipitate cell debris. A glucose standard curve was prepared using Glucose standard (Sigma – G3285) for concentrations of 0 mg/ml, 0.02 mg/mL, 0.04 mg/mL, 0.08 mg/mL and 0.16 mg/mL and a Glycogen standard curve was prepared using Glycogen (Thermo Fisher Scientific, AM9510) for concentrations of 0 mg/ml, 0.02 mg/mL, 0.04 mg/mL, 0.08 mg/mL, 0.16 mg/mL and 0.24mg/mL. Glucose reagent was made by adding 20µL of o-Dianisidine (Sigma F5803) to 1 mL of GO reagent (Sigma, G3660). Glycogen reagent was made by adding 20 μL of Amyloglucosidase (Sigma, A7420) to 1 mL of GO reagent (Sigma, G3660) added with 20µL of o-Dianisidine (Sigma F5803). 30 μL of each standard curve concentration was added into a 96-well plate-reader plate. 6 μL of each sample and 24 μL of 1X PBS were loaded in duplicate to the same 96-well plate-reader plate. 100 μl of Glucose reagent was added to 1 set of samples and the glucose standard curve, 100 μL of Glycogen reagent to the other set of samples and to the glycogen standard curve. Samples were incubated at 37°C for 60 minutes. 100 μl of 12N H2SO4 was added to every well. Absorbance was read at 540 nm (Labtech – SpectroStar Nano).

### Statistical Analysis

Analysis of the images from stained LGs was carried out using a custom written MatLab script (**Ho et al., 2023; available through our lab’s Github account** https://github.com/Tanentzapf-Lab). LGs were individually assessed to count P1-stained cells, total cell numbers based on nuclear staining and the ratio of P1-stained cells to to total cell count was determined. Statistical tests were performed using GraphPad Prism 6 software. Descriptions and results of each test are provided in the figure legends. Sample sizes are indicated in the figure legends. Biological replicates used different larvae. Data for all quantitative experiments were collected on at least three different days, and behavioral experiments were performed with larvae from at least two independent crosses.

## Acknowledgments

We thank the members of the Tanentzapf lab, the TPI lab, the Persing lab and the J. Royet’s lab for the comments on the manuscript. We thank the Bloomington stock center and the Vienna Drosophila resource center for fly stocks. We thank Dr. Istvan Ando for P1 antibodies. We thank Leopold Kurz for his useful guidance.

## Funding

This work was supported by: the Canadian Institutes of Health Research (Project Grant PJT-156277) to GT; by grants from Conseil Régional de Bourgogne-Franche Comté (PARI, ALIMENN grants) to GM; by the CNRS (to PYM, MBG and YG); by Atip-Avenir grant (CNRS Biology, to PYM); by the University of Burgundy Europe (GM and GA); by the GliSFCo grant (ERC, to YG), and Burgundy Council/FEDER (to YG); by the CNRS, ANR BACNEURODRO (ANR-17-CE16-0023-01), Equipe Fondation pour la Recherche Médicale (EQU201603007783) et l’Institut Universitaire de France to J.R. and by the ANR PepNeuron (ANR-21-CE16-0027) to J.R and YG.

## Author contributions

P.Y.M. Conceptualization, Formal analysis, Validation, Investigation, Visualization, Methodology, Writing – original draft, Writing – review and editing; A.N.M. Validation, Investigation, Visualization; D.C. Validation, Investigation; M.B-G and Y.G. Validation, Investigation; G.M. Validation, Investigation; J.R. Validation, Investigation; G.T. Resources, Validation, Writing – original draft, Project administration, Writing – review and editing; R.K. analysis; G.A. Investigation, Visualization; R.M. Validation, Investigation; L.S. Validation, Investigation; E.A. Validation, Investigation; I.C. Validation, Investigation.

## Competing interests

The authors declare no competing interests.

## Data and materials availability

All data are available upon request.

## Supplementary figures

**Figure 1 figure supplement 1.**
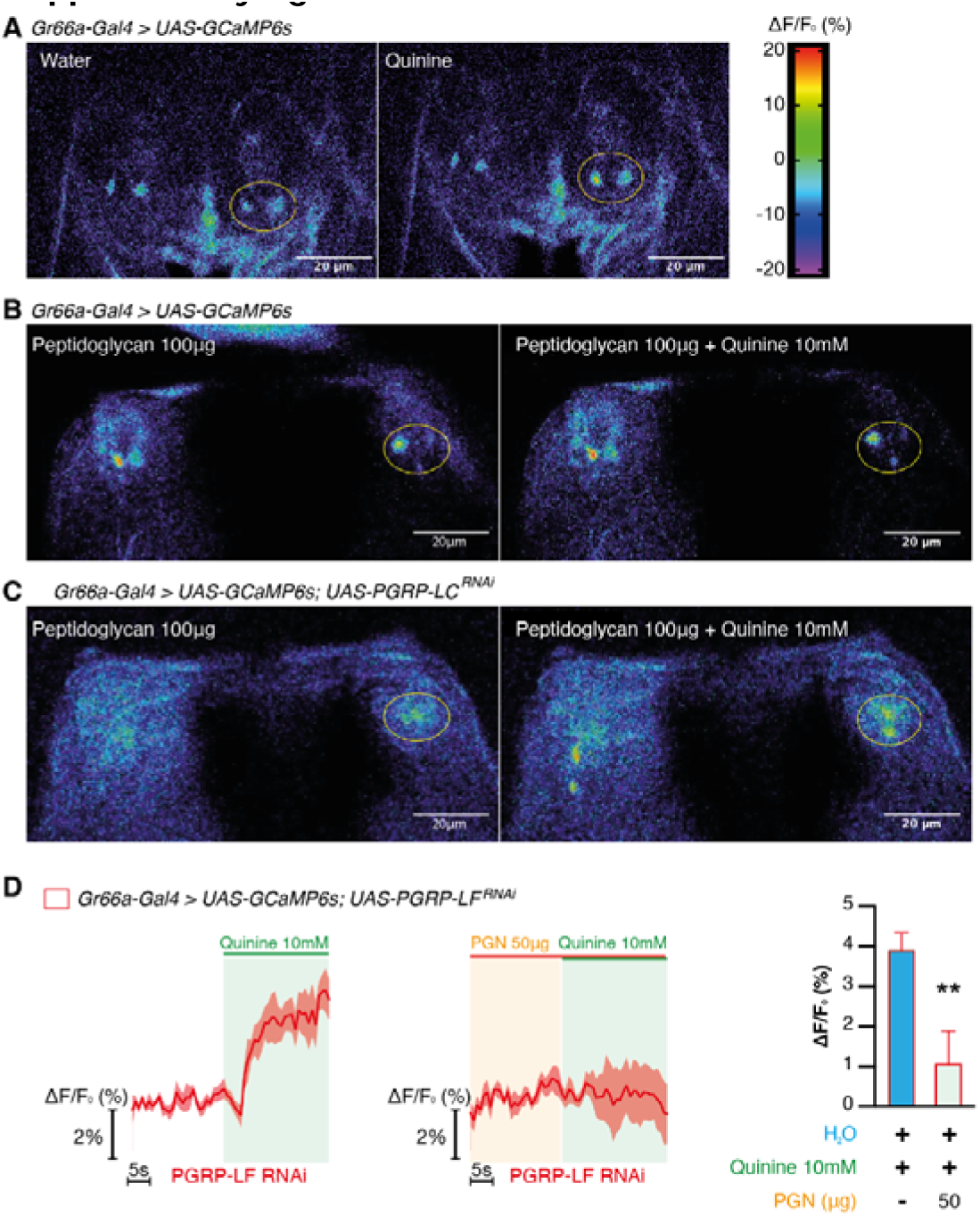
Peptidoglycan sensing inhibits Gr66a taste sensory neurons. (**A**), Representative images showing the GCaMP6s intensity before and after addition of either the control Ringer’s solution (left panels) or the quinine (right panels) in *Gr66a-Gal4>UAS-GCaMP6s* larvae. (**B**), Representative images showing the GCaMP6s intensity before and after addition of either the peptidoglycan solution (left panels) or the quinine+ peptidoglycan (right panels) in *Gr66a-Gal4>UAS-GCaMP6s* larvae. (**C**), Representative images showing the GCaMP6s intensity before and after addition of either the peptidoglycan solution (left panels) or the quinine+ peptidoglycan (right panels) in *Gr66a-Gal4>UAS-GCaMP6s; UAS-PGRP-LC ^RNAi^* larvae. (**D**), Averaged curves of Gr66a neural activity in larvae with PGRP-LF knocked down in response to 10mM quinine preceded by H2O (n = 8) or by peptidoglycan solutions at 50 μg/mL (*n* = 7) (left); and quantification of the responses (right). Values represent mean ± SEM. Statistical tests: Kruskal-Wallis and Dunn’s post hoc; \*\**P* < 0.01.

**Figure 2 figure supplement 1.**
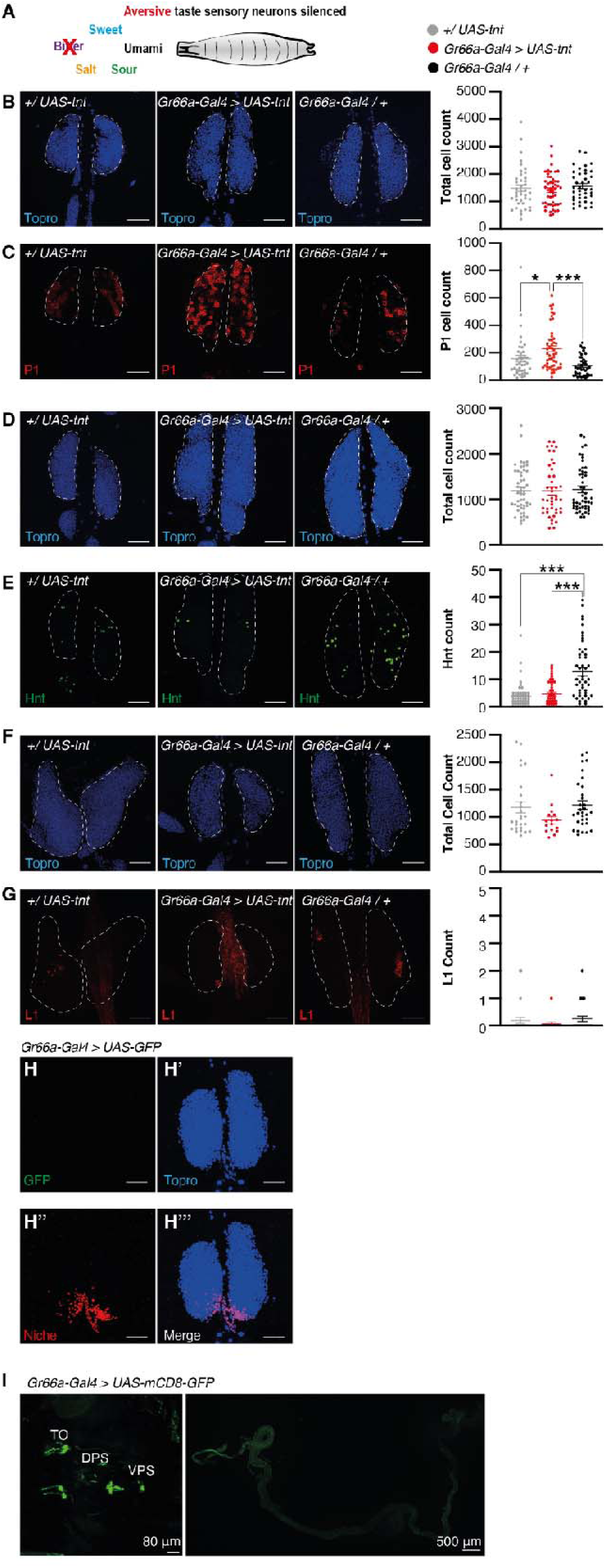
Silencing aversive taste sensory neurons promotes hematopoiesis.) (**A**), Schematic showing taste sensory modalities impaired in *Gr66a-Gal4>UAS-tnt* larvae. (**B**), Staining of lymph gland nuclei (left, Topro-blue) of *+/UAS-tnt* (*n* = 37), *Gr66a-Gal4>UAS-tnt* (*n* = 50), *Gr66a-Gal4/+* flies (*n* = 39), and their quantification (right). (**C**), Staining of lymph gland differentiated cells (left, P1-red) of *+/UAS-tnt, Gr66a-Gal4>UAS-tnt, Gr66a-Gal4 /+* larvae, and their quantification (right). **(D**), Staining of lymph gland nuclei (left, Topro-blue) of *+/UAS-tnt* (*n* = 42), *Gr66a-Gal4 > UAS-tnt* (*n* = 34),*Gr66a-Gal4/+* (*n* = 36) larvae, and their quantification (right). (**E**), Staining of lymph gland differentiated crystal cells (left, Hnt-green) of *+/UAS-tnt* (*n* = 42), *Gr66a-Gal4 > UAS-tnt* (*n* = 34), *Gr66a-Gal4/+* (*n* = 36) larvae, and their quantification (right). (**F**), Staining of lymph gland nuclei (left, Topro-blue) of *+/UAS-tnt* (*n* = 26), *Gr66a-Gal4 > UAS-tnt* (*n* = 16), *Gr66a-Gal4/+* (*n* = 31) larvae, and their quantification (right). (**G**), Staining of lymph gland differentiated lamellocytes cells (left, L1-red) of *+/UAS-tnt* (*n* = 26), *Gr66a-Gal4 > UAS-tnt* (*n* = 16), *Gr66a-Gal4/+* (*n* = 31) larvae, and their quantification (right). (**H**), GFP staining of *Gr66a-Gal4 > UAS-GFP* in the lymph gland (GFP-green), (**H’**), of the nuclei (Topro-blue), (**H’’**), of niche cells (Antp-red). (**H’’’**), Merge of all stainings. (**I**), Staining of Gr66a-Gal4>UAS-GFP larvae mouthpart (left) and guts (right). TO: Terminal Organ, DPS: Dorsal pharyngeal Sensilla, VPS: Dorsal pharyngeal Sensilla. For A to H, scale bars are 50 μm. Values represent mean ± SEM. Statistical tests: one-way ANOVA and Tukey post hoc; \**P* < 0.05; \*\**P* < 0.01; \*\*\**P* < 0.001.

**Figure 2 figure supplement 2.**
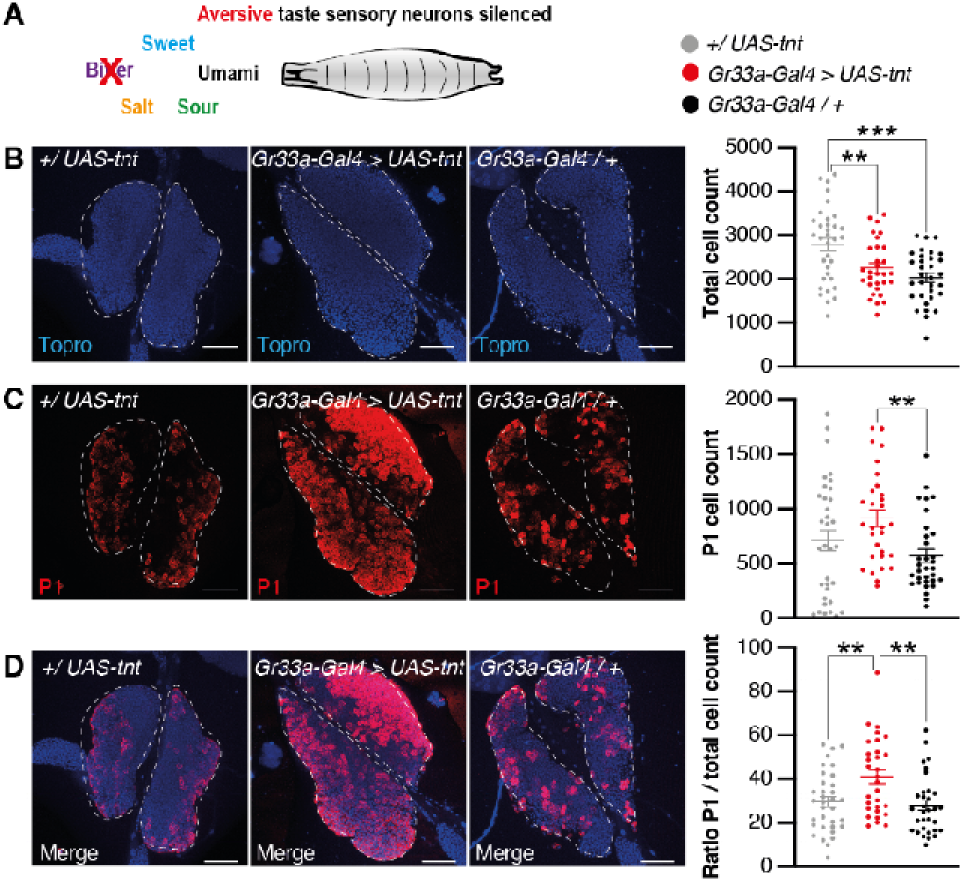
Silencing aversive taste sensory neurons promotes hematopoiesis.) (**A**), Schematic showing taste sensory modalities impaired in *Gr33a-Gal4>UAS-tnt* larvae. (**B**), Staining of lymph gland nuclei (left, Topro-blue) of *+/UAS-tnt* (*n* = 33), *Gr33a-Gal4>UAS-tnt* (*n* = 30), *Gr33a-Gal4/+* flies (*n* = 32), and their quantification (right). (**C**), Staining of lymph gland differentiated cells (left, P1-red) of *+/UAS-tnt, Gr33a-Gal4>UAS-tnt, Gr33a-Gal4 /+* larvae, and their quantification (right). **(D**), Staining of lymph gland nuclei (Topro-blue) and differentiated plasmatocyte cells (left, P1-red) of *+/UAS-tnt*, *Gr33a-Gal4>UAS-tnt*, *Gr33a-Gal4/+* larvae, and quantification of the ratio between the 2 (right). Scale bars are 50 μm. Values represent mean ± SEM. Statistical tests: one-way ANOVA and Tukey post hoc; \**P* < 0.05; \*\**P* < 0.01; \*\*\**P* < 0.001.

**Figure 2 figure supplement 3.**
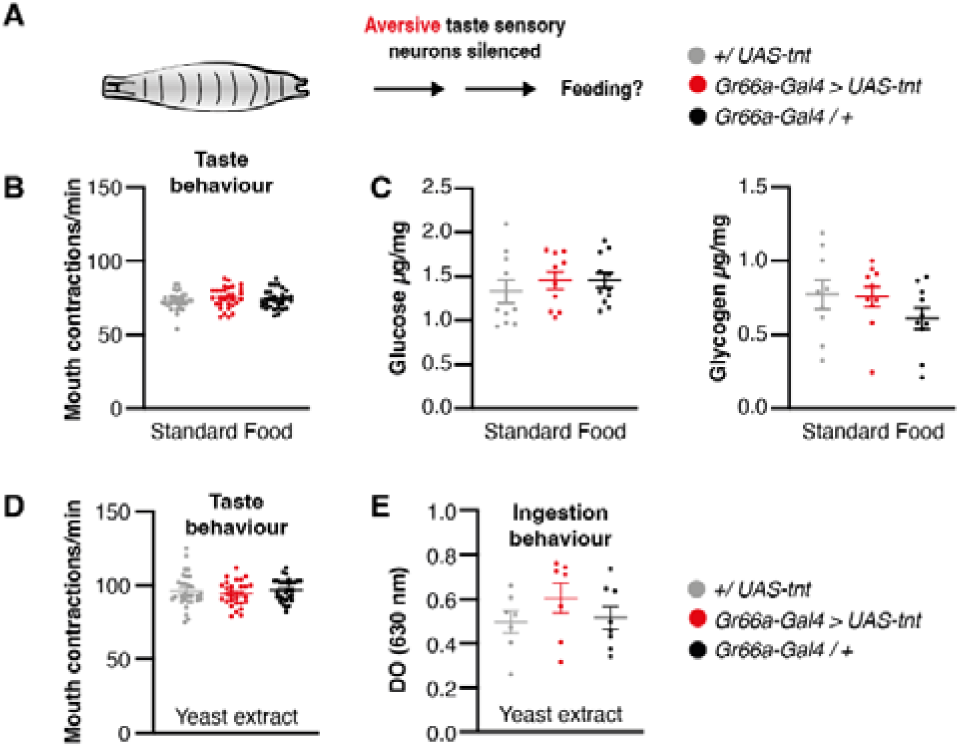
Inhibiting aversive taste neuron activity does not impact feeding. (**A**), Does silencing aversive taste sensing neurons impact feeding? (**B**), Number of mouth contraction per minutes on standard food of *+/UAS-tnt* (*n* = 30), *Gr66a-Gal4>UAS-tnt* (*n* = 30), *Gr66a-Gal4/+* larvae (*n* = 30). (**C**), Glucose (left) and glycogen (right) from *+/UAS-tnt* (*n* = 30), *Gr66a-Gal4>UAS-tnt* (*n* = 30), *Gr66a-Gal4/+* larvae (*n* = 30) raised on standard food. (**D**), Number of mouth contraction (top) per minutes on 2% yeast extract of *+/UAS-tnt* (*n* = 30), *Gr66a-Gal4>UAS-tnt* (*n* = 30), *Gr66a-Gal4/+* larvae (*n* = 30); (**E**), food intake on 2% yeast extract of *+/UAS-tnt* (*n* = 7), *Gr66a-Gal4>UAS-tnt* (*n* = 8), *Gr66a-Gal4/+* larvae (*n* = 7). Values represent mean ± SEM. Statistical tests: one-way ANOVA and Tukey post hoc.

**Figure 2 figure supplement 4.**
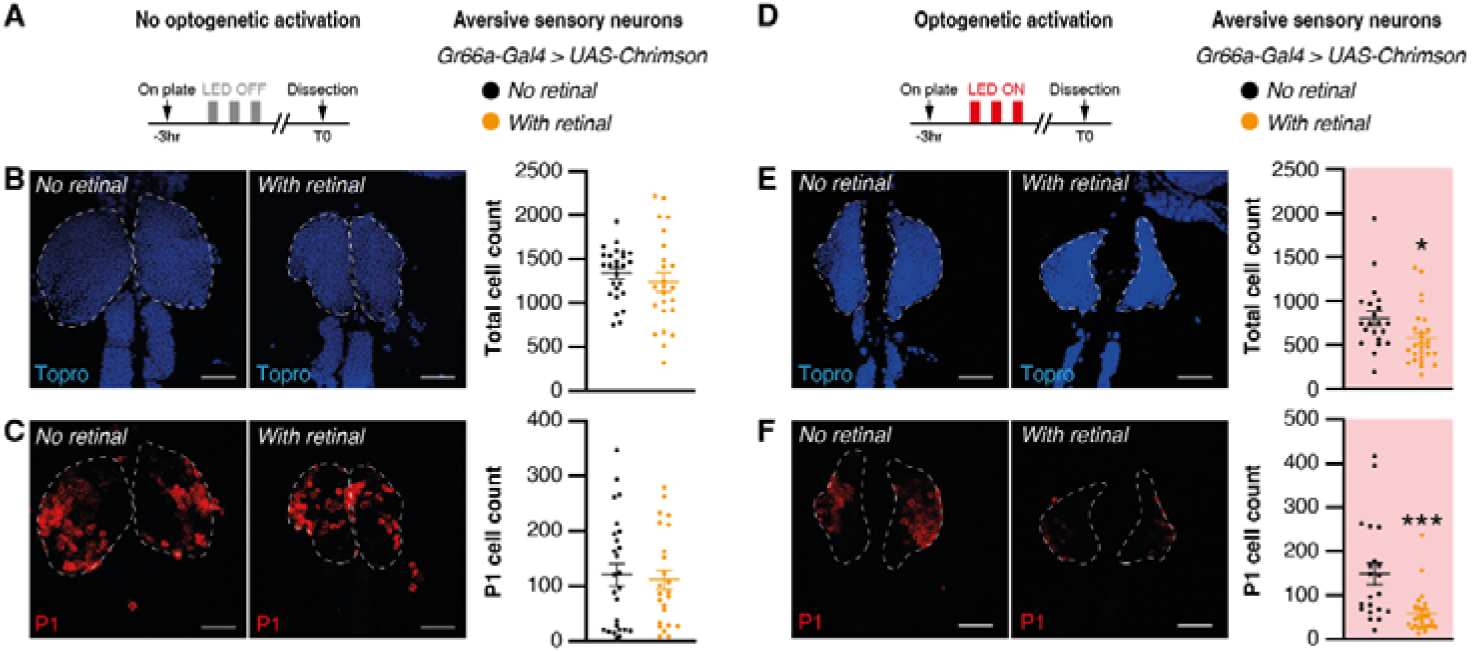
Optogenetic activation of bitter taste sensory neurons inhibits hematopoiesis. (**A**), Schematic showing optogenetic protocol, the OFF light does not activate Gr66a neurons. (**B**), Staining of lymph gland nuclei (left, Topro-blue) and its quantification (right) in non-retinal fed larvae (*n* = 26), and retinal fed larvae (*n* = 24). (**C**), Staining of lymph gland differentiated cells (left, P1-red) and its quantification (right). (**D**), Schematic showing optogenetic protocol, the ON light activates Gr66a neurons in retinal flies. (**E**), Staining of lymph gland nuclei (left, Topro-blue) and its quantification (right) in non-retinal fed larvae (*n* = 18), and retinal fed larvae (*n* = 18). (**F**), Staining of lymph gland differentiated cells (left, P1-red) and its quantification (right). Scale bars are 50 μm. Values represent mean ± SEM. Statistical tests: *t*-test; \**P* < 0.05; \*\*\**P* < 0.001.

**Figure 3 figure supplement 1.**
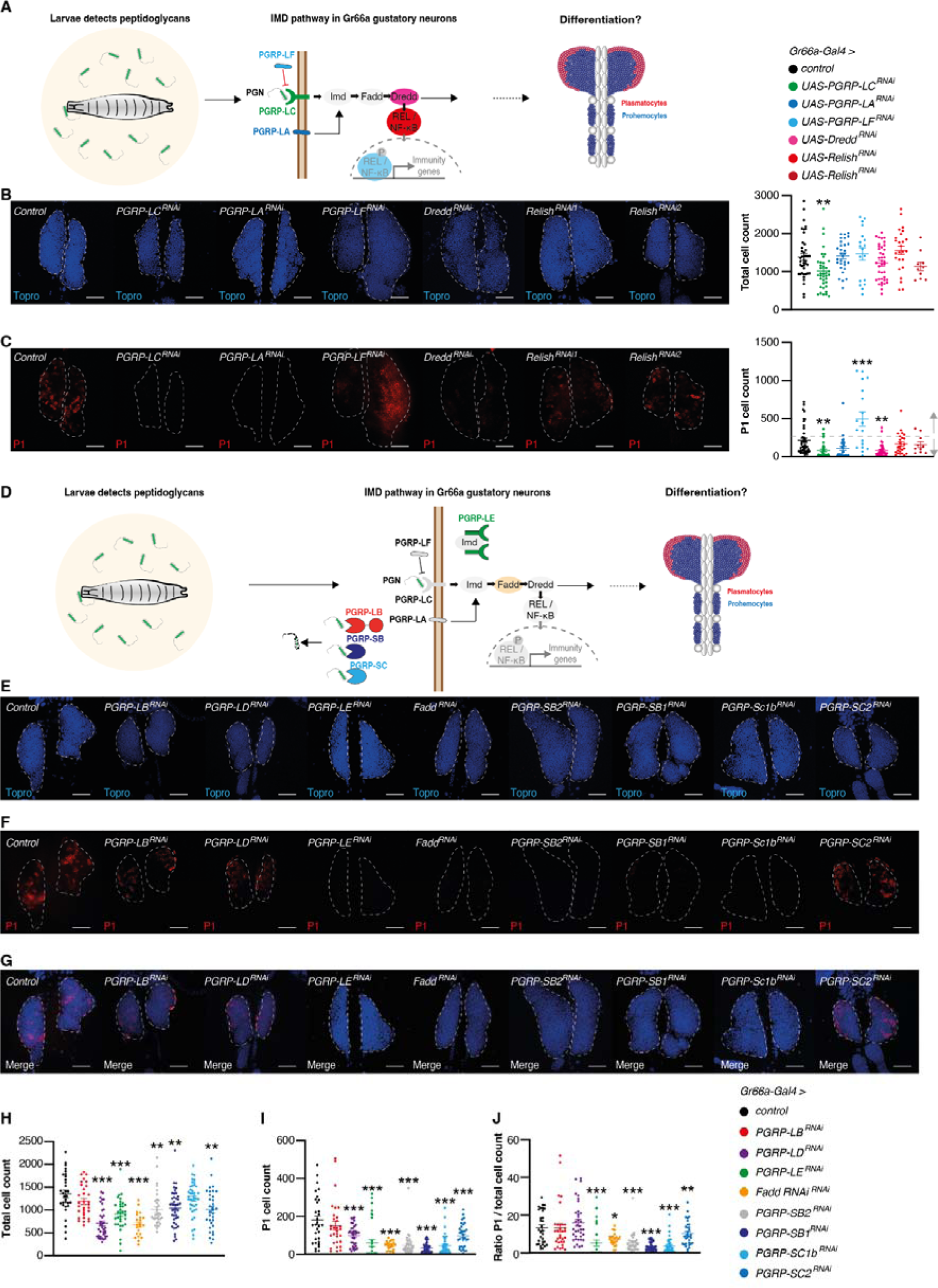
Knocking down PGRPs and Imd pathway in Gr66a taste sensory neurons inhibits hematopoiesis. (**A**), Schematic showing how peptidoglycan detection by the Imd pathway in Gr66a taste sensory neurons could regulate hematopoiesis. (**B**), Staining of lymph gland nuclei (Topro-blue) of *Gr66a-Gal4 > Imd ^RNAi^*larvae (Control, *n* = 42; PGRP-LC, *n* = 38; PGRP-LA, *n* = 30; PGRP-LF, *n* = 18; Dredd, *n* = 39; Relish, *n* = 25, and 10), and their quantification (right). (**C**), Staining of lymph gland differentiated cells (P1-red) of *Gr66a-Gal4 > Imd ^RNAi^* larvae, and their quantification (right). (**D**), Schematic showing how peptidoglycan detection by the Imd pathway in Gr66a taste sensory neurons could regulate hematopoiesis. (**E**), Staining of lymph gland nuclei (Topro-blue) of *Gr66a-Gal4 > Imd ^RNAi^* larvae. (**F**), Staining of lymph gland differentiated cells (P1-red) of *Gr66a-Gal4 > Imd ^RNAi^*larvae. (**G**), Staining of lymph gland nuclei (Topro-blue) and differentiated cells (P1-red) of *Gr66a-Gal4 > Imd ^RNAi^* larvae. (**H**), Quantification of total cell number (Control, *n* = 31; PGRP-LB, *n* = 31; PGRP-LD, *n* = 36; PGRP-LE, *n* = 26; Fadd, *n* = 22; PGRP-SB2, *n* = 31; PGRP-SB1, *n* = 42; PGRP-SC1b, *n* = 41; PGRP-SC2, *n* = 33). (**I**), Quantification of differentiated cells. (**J**), Quantification of the ratio of differentiated cells/total number of cells. (**K**), Schematic showing how Imd, Toll and JNK pathways in Gr66a taste sensory neurons could regulate hematopoiesis. Scale bars are 50 μm. Values represent mean ± SEM. Statistical tests: one-way ANOVA and Tukey post hoc; \**P* < 0.05; \*\**P* < 0.01; \*\*\**P* < 0.001.

**Figure 3 figure supplement 2.**
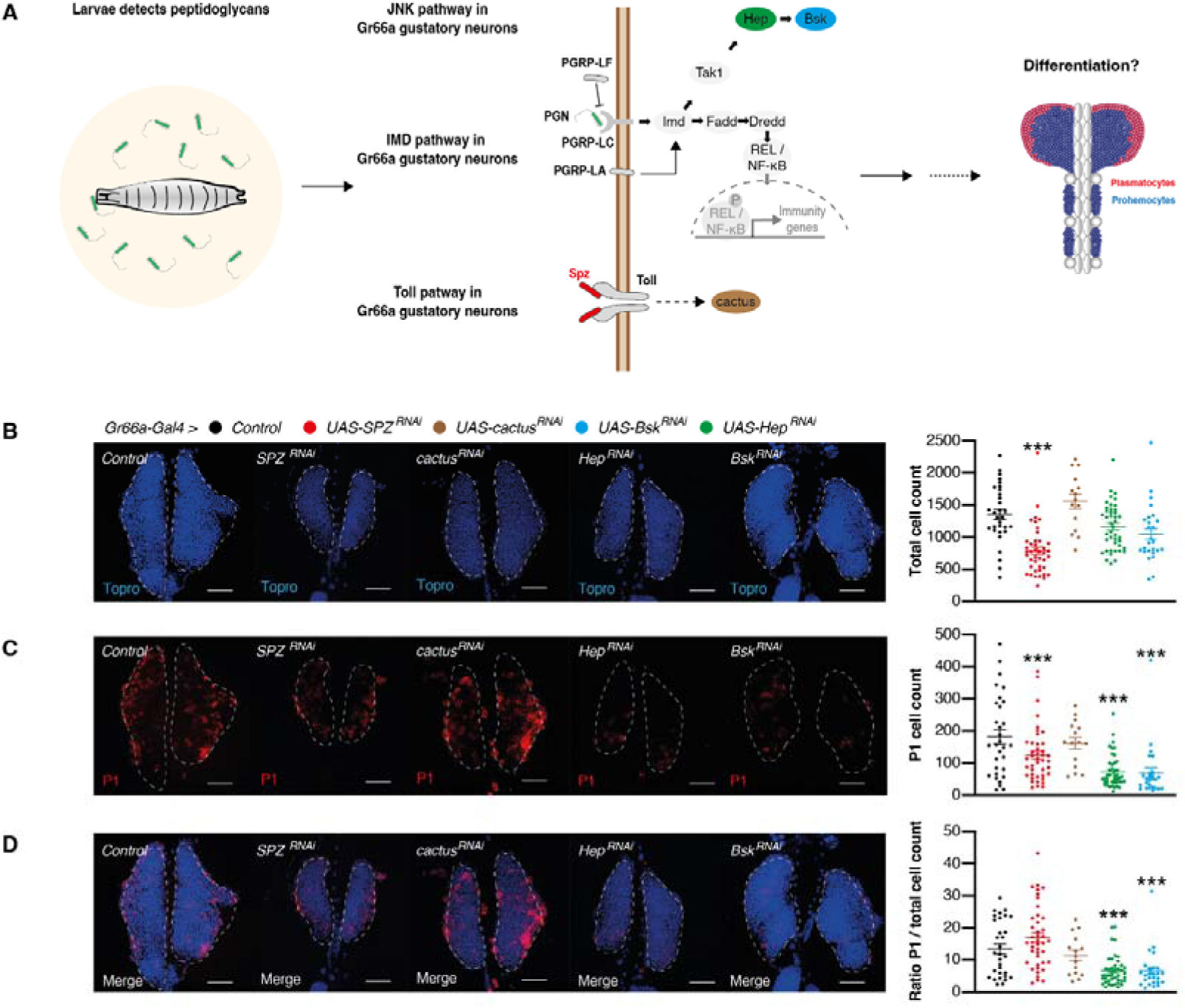
Knocking down JNK pathway in Gr66a taste sensory neurons inhibits hematopoiesis. (**A**), Schematic showing how peptidoglycan detection by the Imd/JNK pathway in Gr66a taste sensory neurons could regulate hematopoiesis. (**B**), Staining of lymph gland nuclei (Topro-blue) of *Gr66a-Gal4 > Toll ^RNAi^* larvae and *JNK ^RNAi^* larvae (Control, *n*= 31; SPZ, *n* = 42; cactus, *n* = 15; Bsk, *n* = 26; Hep, *n* = 43), and their quantification (right). (**C**), Staining of lymph gland differentiated cells (P1-red) of *Gr66a-Gal4 > Toll ^RNAi^* larvae and *JNK ^RNAi^* larvae (Control, *n* = 31; SPZ, *n* = 42; cactus, *n* = 15; Bsk, *n* = 26; Hep, *n* = 43), and their quantification (right). (**D**), Staining of lymph gland nuclei (Topro-blue) and differentiated cells (P1-red) of *Gr66a-Gal4 > Toll ^RNAi^* larvae and *JNK ^RNAi^* larvae (Controls, *n* = 31; SPZ, *n* = 42; cactus, *n* = 15; Bsk, *n* = 26; Hep, *n* = 43), and their quantification (right). Scale bars are 50 μm. Values represent mean ± SEM. Statistical tests: one-way ANOVA and Tukey post hoc; \**P* < 0.05; \*\**P* < 0.01; \*\*\**P* < 0.001.

**Figure 3 figure supplement 3.**
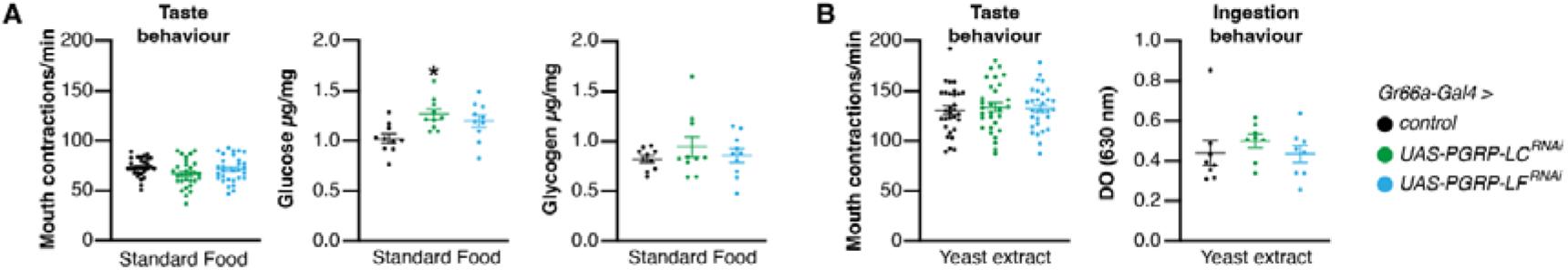
Knocking down PGRPs and Imd pathway in Gr66a taste sensory neurons does not affect feeding behavior. (**A**), Number of mouth contraction per minutes on standard food of *Gr66a-Gal4 > Imd ^RNAi^* larvae (Control, *n* = 30; PGRP-LC, *n* = 30; PGRP-LF, *n* = 30) (left); Glucose (middle) and glycogen (right) from *Gr66a-Gal4 > Imd ^RNAi^*larvae (Controls, *n* =1; PGRP-LC, *n* = 10; PGRP-LF, *n* = 10) raised on standard food. (**B**), Number of mouth contraction per minutes on 2% yeast extract of *Gr66a-Gal4 > Imd ^RNAi^* larvae (Controls, *n* = 30; PGRP-LC, *n* = 30; PGRP-LF, *n* = 30) on standard food (left); food intake on 2% yeast extract of *Gr66a-Gal4 > Imd ^RNAi^* larvae (Controls, *n* = 8; PGRP-LC, *n* = 8; PGRP-LF, *n* = 8). Values represent mean ± SEM. Statistical tests: one-way ANOVA and Tukey post hoc; \**P* < 0.05.

**Figure 3 figure supplement 4.**
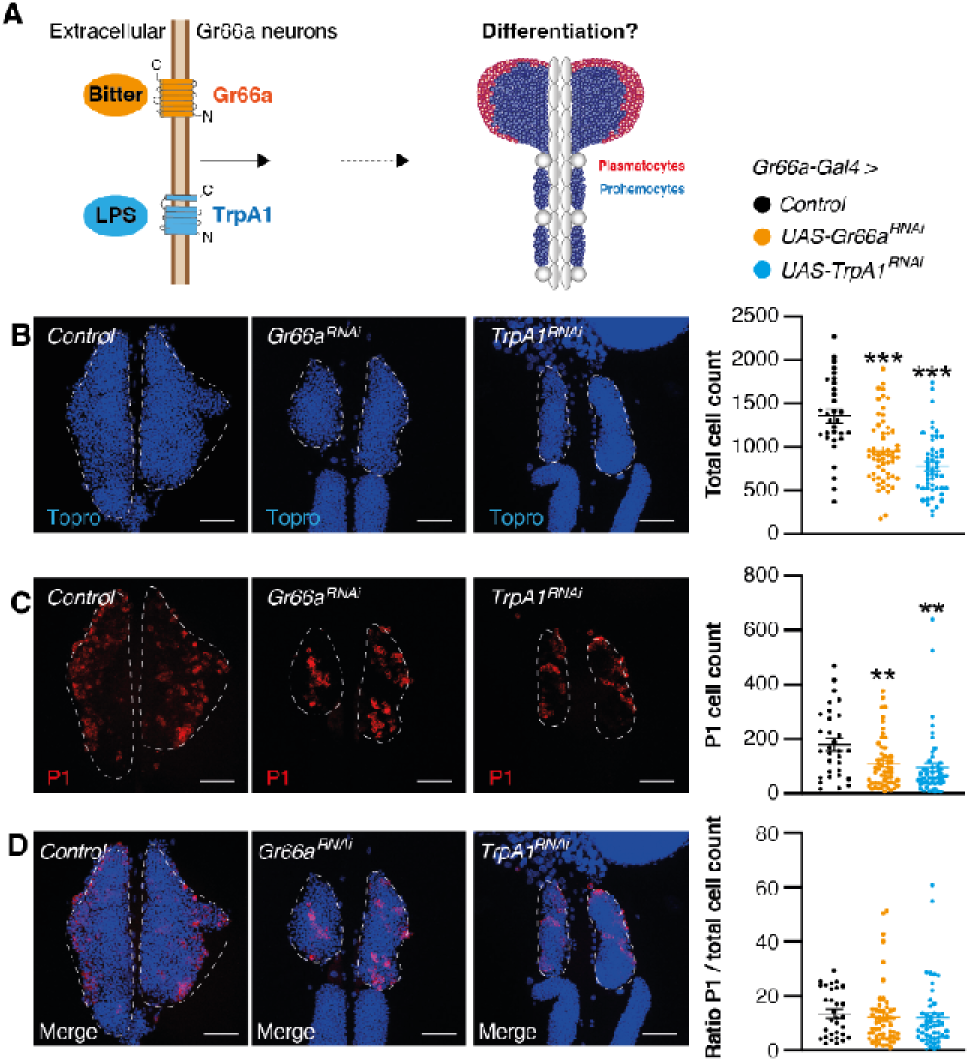
modulating Gr66a and TrPA1 receptors does not modulate hematopoiesis. (**A**), Schematic showing TrpA1 and Gr66a receptor expression in Gr66a taste sensory neurons labeled by Gr66a-Gal4. (**B**), Staining of lymph gland nuclei (left, Topro-blue) *Gr66a-Gal4 > control* (*n* = 31), *UAS-Gr66a^RNAi^*(*n* = 55), and *UAS-TrpA1^RNAi^* (*n* = 49) larvae, and their quantification (right). (**C**), Staining of lymph gland differentiated cells (left, P1-red) of *Gr66a-Gal4 > control*, *UAS-Gr66a^RNAi^*, and *UAS-TrpA1^RNAi^*, and their quantification (right). (**D**), Staining of lymph gland nuclei (Topro-blue) and differentiated cells (left, P1-red) of *Gr66a-Gal4 > control* (*n* = 31), *UAS-Gr66a^RNAi^*(*n* = 55), and *UAS-TrpA1^RNAi^* (*n* = 49) flies, and quantification of the ratio between the 2 (right). Scale bars are 50 μm. Values represent mean ± SEM. Statistical tests: one-way ANOVA and Tukey post hoc; \**P* < 0.05; \*\**P* < 0.01; \*\*\**P* < 0.001.

**Figure 4 figure supplement 1.**
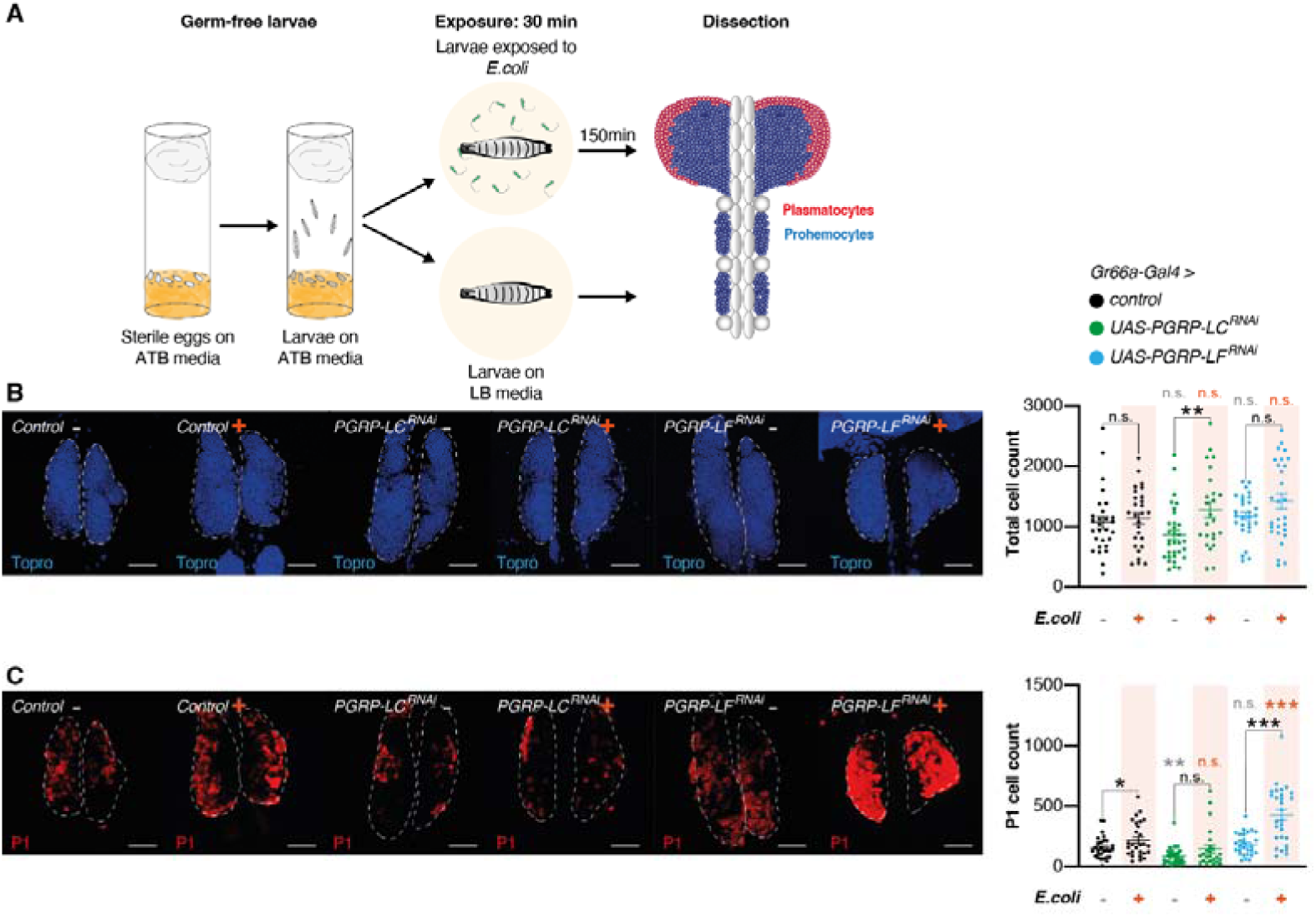
PGRP-LC in Gr66a gustatory neurons mediates increased hematopoiesis after tasting bacteria. (**A**), Schematic representation of the protocol for the exposition of germ-free larvae with *E. coli*. Embryos grow under sterile conditions into a fresh vial containing ATB media. Then larvae are transferred to LB plates containing either *E. coli* or nothing for 30 minutes and placed back to LB media for 150 minutes before dissection. (**B**), Staining of lymph gland nuclei (left, Topro-blue) of *Gr66a-Gal4 > Imd ^RNAi^* flies without exposition to *E. coli* (Control, *n* = 28; PGRP-LC, *n* = 32; PGRP-LF, *n* = 28) and with exposition to *E. coli* (Control, *n* = 27; PGRP-LC, *n* = 24; PGRP-LF, *n* = 28), and quantification (right). (**C**), Staining of lymph differentiated cells (left, P1-red) of *Gr66a-Gal4 > Imd ^RNAi^* flies without exposition to *E. coli* (Control, *n* = 28; PGRP-LC, *n* = 32; PGRP-LF, *n* = 28) and with exposition to *E. coli* (Control, *n* = 27; PGRP-LC, *n* = 24; PGRP-LF, *n* = 28), and quantification (right). Scale bars are 50 μm. Values represent mean ± SEM. Statistical tests: one-way ANOVA, Tukey post hoc, and unpaired *t*-test for comparison of the same genotype between exposed and non-exposed group; \**P* < 0.05; \*\**P* < 0.01; \*\*\**P* < 0.001.

**Figure 5 figure supplement 1.**
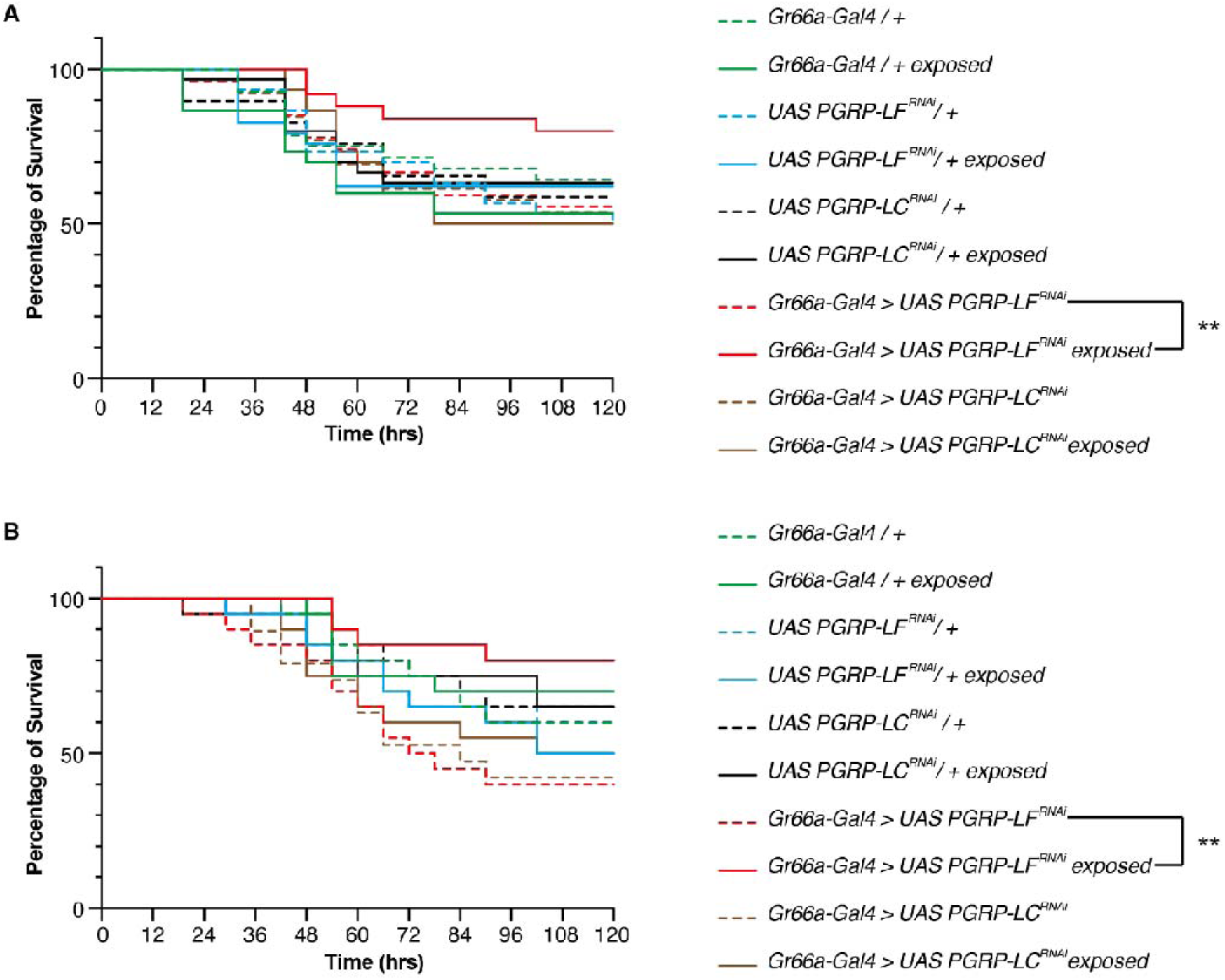
Overactivation of Imd pathway in Gr66a gustatory neurons primes the cellular immune pathway. (**A**), Replicate of the survival experiment: Survival curves following infection with *P.entomophila* of flies previously exposed to *E.coli* (solid lines, *Gr66a-Gal4>6000^TK^*, *n* = 28; *6000^TK^/UAS-PGRP-LF^RNAi^*, *n* = 30; *Gr66a-Gal4>UAS-PGRP-LF^RNAi^*, *n* = 27; *6000^TK^/UAS-PGRP-LC^RNAi^*, *n* = 29; *Gr66a-Gal4>UAS-PGRP-C^RNAi^*, *n* = 26) or to nothing (dashed lines, *Gr66a-Gal4>6000^TK^*, *n* = 30; *6000^TK^/UAS-PGRP-LF^RNAi^*, n = 29; *Gr66a-Gal4>UAS-PGRP-LF^RNAi^*, *n* = 25; *6000^TK^/UAS-PGRP-LC^RNAi^*,*n* = 30; *Gr66a-Gal4>UAS-PGRP-C^RNAi^*, *n* = 30). (**B**), Replicate of the survival experiment: Survival curves following infection with*P.entomophila* of flies previously exposed to *E.coli* (solid lines, *Gr66a-Gal4>6000^TK^*, *n* = 20; *6000^TK^/UAS-PGRP-LF^RNAi^*, *n* = 20; *Gr66a-Gal4>UAS-PGRP-LF^RNAi^*, *n* = 20); *6000^TK^/UAS-PGRP-LC^RNAi^*, *n* = 20; *Gr66a-Gal4>UAS-PGRP-C^RNAi^*, *n* = 20) or to nothing (dashed lines, *Gr66a-Gal4>6000^TK^*, *n* = 20; *6000^TK^/UAS-PGRP-LF^RNAi^*, *n* = 19; *Gr66a-Gal4>UAS-PGRP-LF^RNAi^*, *n* = 20; *6000^TK^/UAS-PGRP-LC^RNAi^*, *n* = 20; *Gr66a-Gal4>UAS-PGRP-C^RNAi^*, *n* = 20). Values represent mean. Statistical tests:one-sided log-rank test; \*\**P* < 0.01.

## Notes

### Competing Interest Statement

The authors have declared no competing interest.

### Summary of Updates

1) Reorganized figures for clarity 2) New experiments added further showing that aversive taste modulates hematopoesis 3) Data consolidate for better clarity

